# Transcriptome-Wide Combinatorial RNA Structure Probing in Living Cells

**DOI:** 10.1101/2020.03.24.006866

**Authors:** Dalen Chan, Chao Feng, Whitney England, Dana Wyman, Ryan A. Flynn, Xiuye Wang, Yongsheng Shi, Ali Mortazavi, Robert C. Spitale

## Abstract

RNA molecules can fold into complex structures and interact with trans-acting factors to control their biology. Recent methods have been focused on developing novel tools to measure RNA structure transcriptome-wide, but their utility to study and predict RNA-protein interactions or RNA processing has been limited thus far. Here, we extend these studies with the first transcriptome-wide mapping method for cataloging RNA solvent accessibility, icLASER. By combining solvent accessibility (icLASER) with RNA flexibility (icSHAPE) data, we efficiently predict RNA-protein interactions transcriptome-wide and catalog RNA polyadenylation sites by RNA structure alone. These studies showcase the power of designing novel chemical approaches to studying RNA biology. Further, our study exemplifies merging complementary methods to measure RNA structure inside cells and its utility for predicting transcriptome-wide interactions that are critical for control of and regulation by RNA structure. We envision such approaches can be applied to studying different cell types or cells under varying conditions, using RNA structure and footprinting to characterize cellular interactions and processing involving RNA.

## Introduction

Precise structural conformations are a hallmark of functional nucleic acid polymers in cells. For example, the genome is arranged in a compact three-dimensional structure, whose dynamics and solvent accessibility are critical for the control of gene expression(1). Methods to measure these structural properties of the genome are now quite mature and can be done genome-wide to understand the interactions between DNA and proteins, and even predict RNA expression outcomes(2). As a corollary, there has also been a more recent emergence of approaches to measure transcriptome-wide RNA structure in cells(3), but these efforts still lag behind in their utility and impact, when compared to the biological significance of DNA-centric measurements discussed above.

The ability of RNA molecules to fold into complex two- and three-dimensional structures is critical for the many biological functions they perform(4, 5). Currently, the majority of reagents used for structure probing can identify base-paired residues within RNA. For example, adenosine and cytosinee residues not involved in Watson-Crick-Franklin pairing can be alkylated by dimethylsulfate(6). Similarly, glyoxals have recently been observed to react with guanosine residues at N-1 and exocyclic N-2 position(7). Selective hydroxyl acylation analyzed by primer extension, or SHAPE, approximates 2’-hydroxyl flexibilities as a proxy for single-strandedness of a residue(8, 9). These long-standing efforts demonstrate the power of regent design to understand and relate chemical reactivity to RNA structure.

Transferring conventional chemical probes from one-RNA-at-a-time measurements to transcriptome-wide scale has proven to be a formidable challenge. Bifunctional RNA structure probing reagents that enable reactivity with RNA and enrichment of reaction sites has been demonstrated to dramatically increase signal-to-noise ratios of RT-stops arising from chemical adducts(10, 11). We previously developed icSHAPE, which features a bifunctional chemical probe to measure nucleobase flexibility through RNA hydroxyl acylation transcriptome wide(10) (**Fig. 1, a**). Solvent accessibility, an orthogonal property of RNA, has been more difficult to probe inside cells, requiring the use of a synchrotron radiation light source(12). We recently described the development of LASER, or <u>L</u>ight <u>A</u>ctivated <u>S</u>tructural <u>E</u>xamination of <u>R</u>NA(13). LASER relies on the facile light activation of a nicotinoyl azide to a nitrenium ion, which undergoes electrophilic aromatic substitution with electron rich purine residues at their C-8 nucleobase position. While this initial work demonstrated the utility of LASER to read out solvent accessibility, it still remained a critical challenge to extend these efforts to a transcriptome-wide scale and evaluate this approach to understand RNA structure and function more broadly.

**Figure 1:**
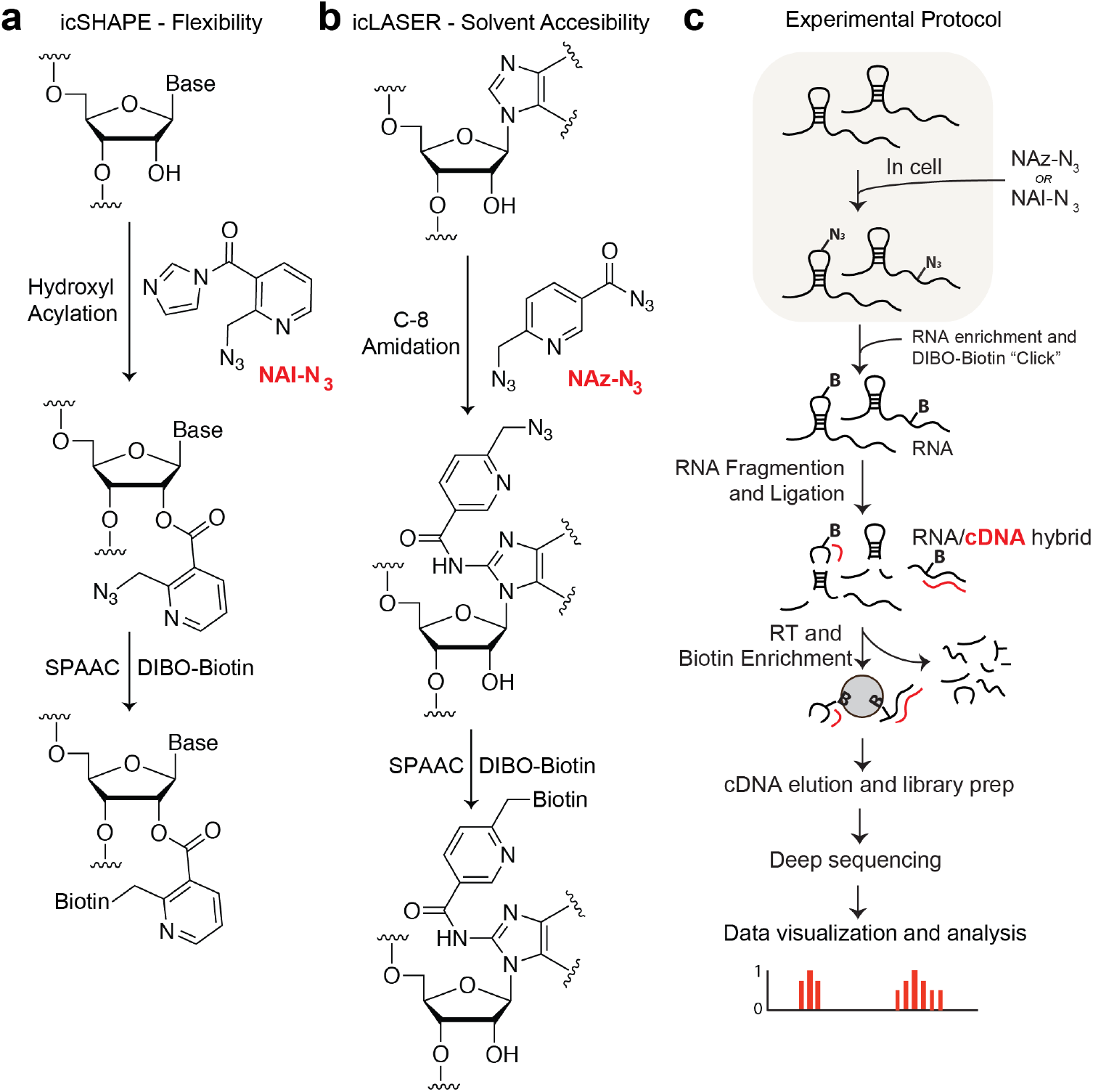
Schematic for icSHAPE and icLASER. **a.** Chemical protocol for icSHAPE(10). **b.** Chemical protocol for icLASER. **c.** Protocol for library preparation for icSHAPE and icLASER, which could be integrated in the same protocol method for RNA structure probing, transcriptome-wide.

Herein we describe our efforts to greatly expand the toolbox of RNA structure probing reagents, and demonstrate their utility to understand RNA structure in cells as well as interactions involving RNA to control RNA processing and RNA-protein interactions. We designed, synthesized, and implemented a bifunctional LASER probe to measure transcriptome-wide purine C-8 solvent accessibility (**Fig. 1, b,** *in vivo* click LASER, icLASER). Our design employed an alky azide functional group for enrichment, therefore permitting a common biochemical and computational approach to identify RNA-reagent adducts for measuring RNA structure (**Fig. 1, c**). By implementing this novel bi-functional probe, icLASER provides the first transcriptome-wide measurement of solvent accessibility. Towards the goal of integrating multiple, orthogonal measurements of RNA structure, we develop an approach to directly compare icSHAPE (hydroxyl acylation; flexibility) and icLASER (solvent accessibility). We highlight the power these data and this strategy in the context of predicting RNA-protein interactions and RNA polyadenylation. Our results demonstrate that combinatorial RNA structure probing can be employed to better understand RNA structure and processing in cells transcriptome-wide, and is thus a highly-useful complement and/or predictor of critical interactions that control RNA biology.

## Results

### Development of a bifunctional LASER probe

An optimal probe for icLASER would retain the selectivity of NAz and also allow for enrichment using conditions non-harmful to RNA. As such, NAz-N_3_, which we predicted would be able to undergo light transformation to the activated nitrenium ion, would also preserve the alkyl azide for Cu-free “click” after RNA adduct formation. Notably, aroyl azides are amenable to activation by lower energy long wavelength light, whereas alkyl azides need higher energy short wavelength light for activation(14–16). We hypothesized these special characteristics of azide stability would permit photo-specific activation of NAz-N_3_ for icLASER probing (**Fig. 2, a**).

**Figure 2:**
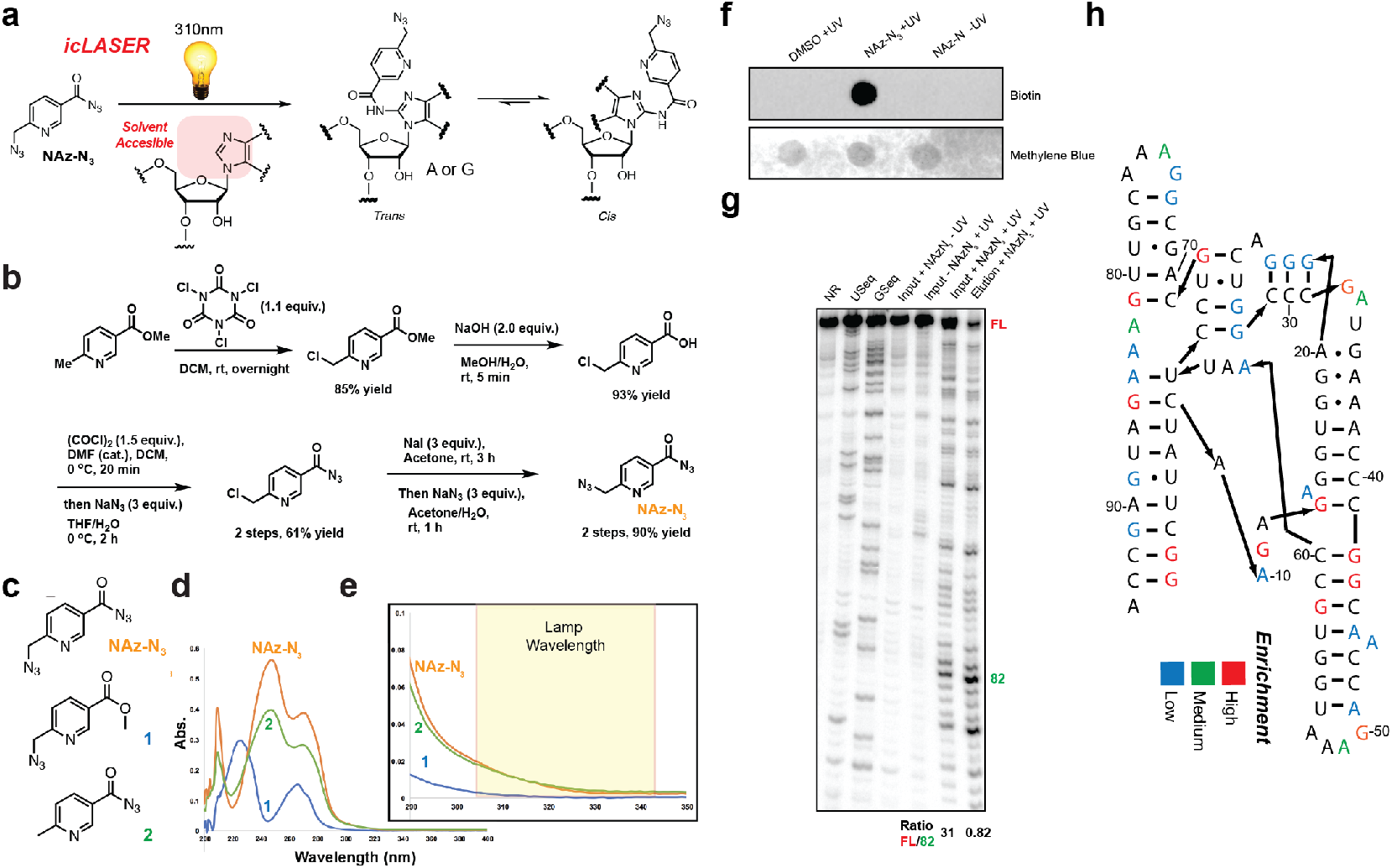
Development of a bi-functional probe for icLASER. **a.** Reaction schematic for LASER, with C-8 amidation utilizing NAz. **b.** Synthetic scheme for the probe NAz-N_3_. **c.** Structure of NAz-N_3_ and control probes for analysis of UV-VIS spectroscopy and light activation. **d.** UV-VIS spectra of compounds from panel c. **e.** Zoomed in UV-Vis spectra and corresponding wavelength used for light-activation of NAz-N_3_ and control compounds in panel c. **f.** Streptavin-HRP dot blot of NAz-N_3_ – modified RNA. RNA was incubated with NAz-N_3_ in the presence of long-wavelength UV light. RNA was precipitated and conjugated with biotin using SPAAC, as denoted in Fig. 1. Followng SPAAC, RNA was blotted. **g.** Denaturing gel electrophoresis of modified SAM-I control RNA with NAz or NAz-N_3_. RNA was incubated with NAz-N_3_ in the presence of long-wavelength UV light. RNA was precipitated and conjugated with biotin using SPAAC. RNA was then enriched over magnetic streptavidin beads and eluted. Eluted RNA was reverse transcribed with ^32^P labeled primer and cDNA analyzed on denaturing gel. Ratio of full length cDNA to modification was calculated against position 82 in the denaturing gel to demonstrate de-enrichment of the full length in comparison to the enriched modified position. **h.** Analysis of enriched positions from panel f.

To test our hypotheses, we synthesized NAz-N_3_ and compounds 1 and 2 (**Fig. 2, b; Supplementary Information**). The NAz-N_3_ probe is synthesized efficiently from commercially available methyl 6-methylnicotinate in six steps in 43% overall yield. Treatment of methyl 6-methylnicotinate with trichloroisocyanuric acid generated monochloro substitution on the methyl group selectively, followed by alkaline hydrolysis of the ester to afford the corresponding carboxylic acid. The acetyl azide is installed routinely by transformation of the carboxylic acid into acetyl chloride using oxalyl chloride and azide substitution. The above monochloro substituent is finally converted to iodide through a Finkelstein reaction, which is replaced subsequently by azide to furnish NAz-N_3_. Synthesis of 1 and 2 is reported in **Supplementary Information**.

We obtained UV-Vis spectra of these compounds, to understand differences in their absorbance properties (**Fig. 2, c**), and observed that only NAz-N_3_ and **2** have absorption in the long wavelength UV region used previously by us in LASER and known to be specific for aroyl azides (**Fig. 2, d&e**)(13). Consistent with what others have observed with azide-specific activation, these data suggest that light activation of NAz-N_3_ should be specific to the aroyl azide. To test reactivity and preservation of a SPAAC-compatible azide, we incubated an *in vitro* transcribed SAM-I riboswitch with NAz-N_3_ and exposed the solution to (+/-) long wavelength UV light. Following irradiation, we isolated the RNA and incubated the RNA with dibenzylcyclooctyne-conjugated to biotin (DBCO-biotin). We observed biotin conjugation on a streptavidin dot blot only in the sample incubated with long wavelength UV light and NAz-N_3_ (**Fig. 2, f**).

We next tested if NAz-N_3_ enriched RNA would produce similar truncation profiles to NAz probing (LASER). We used NAz-N_3_ modified riboswitch RNA, biotinylated the RNA using SPAAC, and subjected it to streptavidin-coated magnetic bead enrichment. Biotinylated RNA was then eluted and compared against input RNA. Each sample was then reverse transcribed with a ^32^P-cDNA primer and analyzed by denaturing gel electrophoresis. As shown in **Fig. 2, g,** the NAz-N_3_ enriched samples displayed significantly higher signal for structure stop cDNA and dramatically reduced full-length extensions. Finally, NAz-N_3_ elution mapped onto sites of modification similar observed with NAz (A ɖ G residues; **Fig. 2, h**). Further, we also demonstrated that NAz-N_3_ can modify RNA and measure RNA structure in cells, and observed differences *in vitro* and in cells that correspond to cDNA profiles to that of the parent compound NAz(17) (**Supplementary Figure 1**). Overall, this result demonstrates that NAz-N_3_ is capable of enrichment using SPAAC reactions, yet still yields consistent truncation signatures for structural analysis.

### Transcriptome-wide implementation of NAz-N_3_

We modified K562 cells (*in vivo*) and K562 total RNA isolated and re-folded outside of cells (*in vitro)* in independent experiments with NAz-N_3_ (icLASER) and NAI-N3 (icSHAPE). We applied our well-established protocol for the isolation of poly-adenylated (polyA) RNA and mapping of biotin-conjugated RNA structure probes through deep sequencing (**Figure 1, Supplementary Methods)** (18, 19), which generated more than 100 million reads for each dataset (**Supplementary Figure 2**). Each dataset had strong agreement with each other and RT stops were enriched for G and A.

Mapping RT stops from icLASER enriched libraries revealed a marked enrichment of A and G residues, which was expected given the reactivity profile of the NAz-N_3_ probe (**Supplementary Figures 2 & 3**). This profile was unique to icLASER, whereas for icSHAPE we observed a slight enrichment for adenosine stops, as seen previously (**Supplementary Figure 3**) (10). Importantly, we find a strong correlation between +UV samples, whereas, the distribution of +UV and -UV RT-stops showed very low correlation (**Supplementary Figure 4**). Comparison of the icSHAPE and icLASER stops showed strong intra-reagent correlation but weaker inter-reagent correlation, suggesting each probe measured a discrete set of positions in the transcriptome.

As an initial quality control, we analyzed the icLASER and icSHAPE data against manual footprinting on an RNA structure in which we had reasonably high-resolution data: the ribosome. We focused on a section of 18S rRNA has been previously interrogated by footprinting(13) (**Fig. 3, a & b**). The manual footprinting for NAI-N3 in this region showed some slight differences between in and outside of cells (**Fig. 3, c**). In similar fashion, our icSHAPE data had similar overall reactivity between the two experimental setups (**Fig. 3, d**). Analysis of the RNA in this region (Fig. 3, e) revealed that it was solvent protected by rRNA and ribosome binding proteins, but was largely still single stranded and thus would be reactive toward SHAPE reagents. We also performed comparison probing with NAz-N_3_ on the same stretch of rRNA and observed marked differences between in and outside the cells (**Fig. 3, f**). Consistent with this analysis was the reactivity profile for icLASER (**Fig. 3, g**), which is supported by the solvent protected nature of this stretch of RNA in the 18S ribosome CryoEM model (**Fig. 3, h**). Overall, these results suggest that icLASER and icSHAPE are both measuring environment-unique aspects of RNA structure in and outside cells.

**Figure 3:**
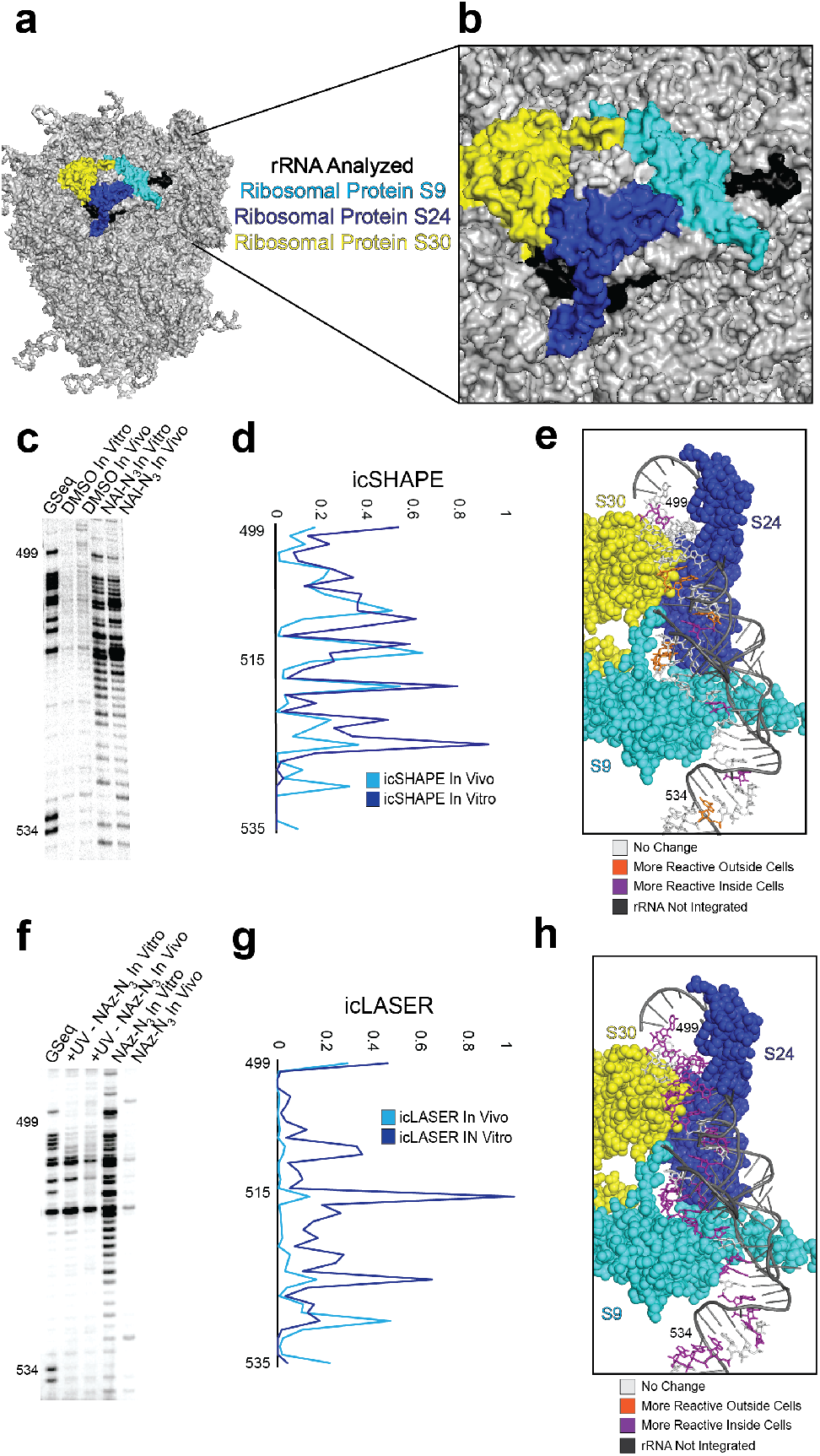
Analysis of icLASER and icSHAPE RT-stops. **a.** CryoEM model of the 80S ribosome (PDB 4v6x). **b.** Zoom in of rRNA section interrogated in for comparison footprinting. **c.** Denaturing gel analysis of NAI-N3 reactivity. In vitro re-folded RNA or cells were subjected to incubation with NAI-N3. Modification sites were analyzed by reverse transcription. **d.** icSHAPE reactivity profile over the same 18S rRNA region as in panel c. **e.** Structural analysis of 18S rRNA with differential reactivity from icSHAPE colored. **f.** Denaturing gel analysis of NAz-N_3_ reactivity. In vitro re-folded RNA or cells were subjected to incubation with NAz-N_3_. Modification sites were analyzed by reverse transcription. **g.** icLASER reactivity profile over the same 18S rRNA region as in panel f. **h.** Structural analysis of 18SrRNA with differential reactivity from icLASER colored.

As this is the first transcriptome-wide map of solvent accessibility probing with icLASER, we also analyzed mRNA profiles. We observed that start codons are largely open and solvent accessible with high reactivity at the A and G of the AUG (**Supplementary Figure 5**). The stop codon also displayed high reactivity in the last two positions (**Supplementary Figure 5**), and both of these observations are in line with previous measurements using icSHAPE and DMS-seq (20).To understand what regions of icLASER and icSHAPE signal could be due to conditions specific to living cells, we compared the *in vivo* and *in vitro* profiles as previously described for icSHAPE (VTD = *in vivo* profile – *in vitro* profile, (21)). Specifically, we examined all hexamers across the transcriptome, and consistent with our previous reports, icSHAPE structure was overall higher *in vivo* (**Supplementary Figure 6.**). This is hypothesized to be correlated with structural remodeling *in vivo,* in comparison to *in vitro* (10). In contrast, we observed a bimodal distribution of solvent accessibility differences in and outside of cells with icLASER. Hexamers that include the Kozak sequence showed small VTD differences, which is in line with previous reports with icSHAPE(21). However, for icLASER, hexamers containing RNA binding protein motifs had lower maximal averaged VTD values. This observation may be consistent with the known reactivity of LASER probing, which can result in a change in chemical reactivity in cells due to protein footprinting, suggesting that icLASER could be used to map protein-RNA interactions. Overall, these results demonstrate the robustness of icLASER probing and demonstrate its utility in mapping the structures of RNAs in and outside of cells.

### Global prediction of protein footprints by combining structure probing tools

Despite the importance of RNA-protein interactions and the interplay of RNA structure and protein binding, there has thus far been a lack of transcriptome-wide analysis of RNA structure motifs with the express goal of attempting to predict binding sites. To address this gap, we assessed if icLASER, icSHAPE, or their combination could be used to predict RNA-protein interactions (**Fig. 4**). As a reference for experimentally measured RNA binding protein (RBP) sites, we leveraged enhanced individual nucleotide resolution CLIP (eCLIP) datasets from ENCODE collected from K562 cells (22, 23). We focused on RBPs that had significant binding in 3’-UTRs (ensuring high sequence coverage in icLASER and icSHAPE data), binding motifs with at least three A or G residues, targets expressed at greater than or equal to 5 RPKM. As control regions, we sampled the same number of sites with matching RNA motifs not called as peaks in the eCLIP data.

**Figure 4:**
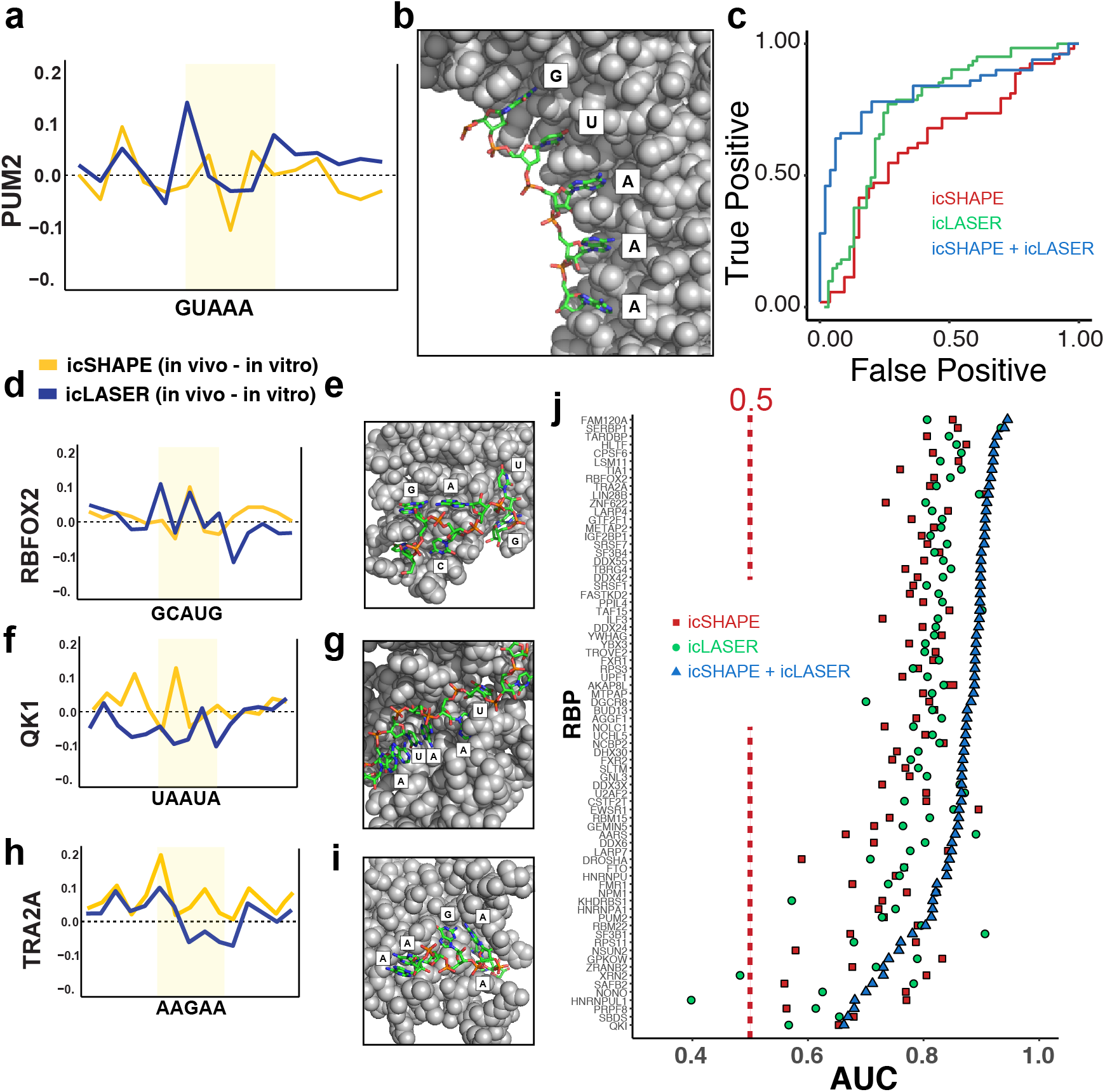
Utilizing icSHAPE and icLASER data to predict RNA-protein interactions transcriptome-wide. **a.** icSHAPE and icLASER difference maps between RNA probed inside living cells and in vitro centered at the motif for PUM2. **b.** X-Ray structure of PUM2 bound to RNA (PDB 3Q0Q). **c.** ROC analysis for predicting RNA-protein interactions for PUM2 eCLIP sites. **d.** icSHAPE and icLASER difference maps between RNA probed inside living cells and in vitro centered at the motif for RBFOX2. **e.** X-Ray structure of RBFOX2 bound to RNA (PDB 2ERR). **f.** icSHAPE and icLASER difference maps between RNA probed inside living cells and in vitro centered at the motif for QK1. **g.** X-Ray structure of QK1 bound to RNA (PDB 4JVH). **h.** icSHAPE and icLASER difference maps between RNA probed inside living cells and in vitro centered at the motif for TRA2A. **i.** X-Ray structure of TRA2A bound to RNA (PDB 2KXN). **j.** ROC analysis for predicting RNA-protein interactions using icLASER and/or icSHAPE structure probing. For each RNA-binding protein, we selected eCLIP bound sites *in vivo* and *in vitro.* A portion of this dataset was used as a training set, and the remainder was used to test the classifier. The classifier was trained using icSHAPE profiles, icLASER profiles, or both.

We first focused on PUM2, a well-established RNA binding protein that is important in regulating RNA localization and translation, and has high-resolution structures for comparison (**Fig. 4, a**)(24). Comparative icLASER and icSHAPE profiles in and outside of cells yielded key differences that likely reflect protein binding (**Fig. 4, b**). First, the G1 position has much higher icLASER signal *in vivo* and this position has its C-8 atom pointed directly out into solvent, suggesting that the protein may be holding this portion of the RNA in this conformation. Positions A3-5 have minimal differences *in vivo* and *in vitro* by icLASER and these positions are also out away from the structure. In contrast, the icSHAPE profile has lower reactivity at position A3, and this may be due to different levels of stability from binding. These differences overall, and the reactivity differences observed at RNA binding sites overall (**Supplementary Figure 6**), suggested to us that structure probing using these reagents could be utilized to predict RNA-protein interactions. To test this, we employed support vector machine learning (SVM)(10), training the classifier using the VTD profiles at eCLIP-defined binding sites *in vitro* and *in vivo.* We trained the classifier using icSHAPE, icLASER, or combined profiles, using randomly selected sites as a training set. We evaluated performance using the area under the reporter operator characteristic (ROC) curve for PUM2 and observed that combining icSHAPE and icLASER profiled dramatically increased the ROC area and predictability. (**Fig. 4, c)**. These results suggest that a larger analysis of comparative structure probing could enable more analyses and prediction of RNA-protein interactions in cells.

We expanded our initial analysis to other RBPs that have reported crystal or NMR structures bound to cognate RNA sequences. RBFOX2 (**Fig. 4, d&e**) displayed a similar icSHAPE structure profile to our previous efforts with icSHAPE in mouse embryonic stem cells(21). icLASER profile differences showed high reactivity at the G1 position, and correspondingly the C-8 of G1 is pointed out to solvent in the crystal structure. The same is true for A3, where the C-8 position is solvent exposed. Quaking1 (QK1) (**Fig. 4, f&g**) shows generally lower icSHAPE reactivity across its binding motif. These positions are also protected from solvent either through direct protein interactions or shielding from phosphate backbone, as is the case for A3 in the motif. Lastly, TRA2A (**Fig. 4, h&i**) displayed correlation at positions 1 and 2 (AA); these are both solvent exposed and likely to be highly dynamic, and as such had high icLASER and icSHAPE reactivity. The last two A residues were protected from solvent by protein interactions and would likely be stabilized to limit their SHAPE reactivity. Overall, these data show icLASER and icSHAPE provide non-overlapping nucleotide resolution at RBP binding sites that can read out the biophysical conformation of the bound RNA in a manner that corresponds to each method’s chemical properties.

To further expand our analysis of combined chemical probing we evaluated performance using the area under the reporter operator characteristic (ROC) curve for each protein (**Fig. 4j**). Using this approach, icLASER and icSHAPE data alone were able to predict between 50-70% of the binding sites for 75 RBPs analyzed herein. Overall, the predictability was higher for icLASER in comparison to icSHAPE. However, when we combined the icLASER and icSHAPE datasets, we observed a dramatic increase in predictive power for protein binding sites and were now able to predict over 85% protein occupancy on RNAs. We suspect the robust, but in some instances imperfect, prediction may be due to the inability of LASER probing to differentiate RNA protection from proteins versus RNA tertiary structure. It is worth noting that for 75 RBPs (300 experiments: biological duplicate eCLIP + biological duplicate background) we are predicting their binding with 8 experiments (icSHAPE/icLASER biological duplicate + biological duplicate background). Overall, these results support the notions that: (1) RNA solvent accessibility probing can be utilized to predict RNA-protein interactions, (2) icLASER data can be utilized in conjunction with CLIP datasets to further support protein occupancies determined by orthogonal methods, and (3) by combining icLASER and icSHAPE, robust and transcriptome-wide predictions of many RBPs is possible without protein-centric techniques such as CLIP.

### Predicting polyA sites in RNA with structure probing

Analysis discussed previously (in relation to VTD; **Supplementary Figure 6**) revealed hexameric sequences that have differences inside and outside of cells. One such sequence that had higher in-cell reactivity for icLASER (and icSHAPE) was the polyadenylation signal sequence (PAS) within the 3’-UTRs(25). Given the measurable VTD value, we explored if our structure probing data could be used predict polyadenylation transcriptome-wide. This would be an extremely important further validation of our approach to use structure probing for predicting RNA-protein interactions and potential processing events.

To experimentally determine PAS sites, for SVM analysis, we generated the first polyadenylation sequencing data (PAS-seq (26)) for K562 cells. PAS-seq uses polyA tail priming to identify the sites of polyA tail selection (**Fig. 5, a**). Inspection of PAS reads demonstrated clear read buildup at the 3’-end of transcripts (**Fig. 5, b**) and we obtained sites of high and low PAS read depth (**Fig. 5, c**). We then compared the icLASER and icSHAPE profiles at the PAS sites and sequences with the same motif which are not annotated to be PAS sequences (negative control) (27). We noticed a striking structural difference between them: a large peak at the first two adenosine residues, followed by a drop in icLASER signal for the remaining UAAA (**Fig. 5, d**). To understand if these differences were related to a biophysical conformation of the RNA in an active PAS-recognition complex, we examined a newly published structure(28). This structure contains three large proteins which form into a complex and recognize the AAUAAA motif (**Fig. 5, e**). Close inspection of the structure revealed that the two five prime adenosine residues are completely solvent exposed, whereas the UAAA residues are solvent protected by protein binding. Impressively, the two AA residues have their C-8 positions (site of icLASER reaction) completely exposed (**Fig. 5, e**). Consistent with our icLASER data, sites with high icLASER signal at the first two adenosine residues had very high coverage by PAS-seq. This data further suggested to us that icLASER signal alone could be used to predict polyA site selection. To test this hypothesis, we utilized SVM(10) and demonstrated that icLASER and icSHAPE used together had an AUC of 0.9 for predicting PAS sites (**Fig. 5, f**). These data nicely demonstrate that icLASER (and combined icLASER and icSHAPE) can be used to predict sites of posttranscriptional regulation and could be integrated with orthogonal datasets to interpret posttranscriptional processing of RNAs.

**Figure 5:**
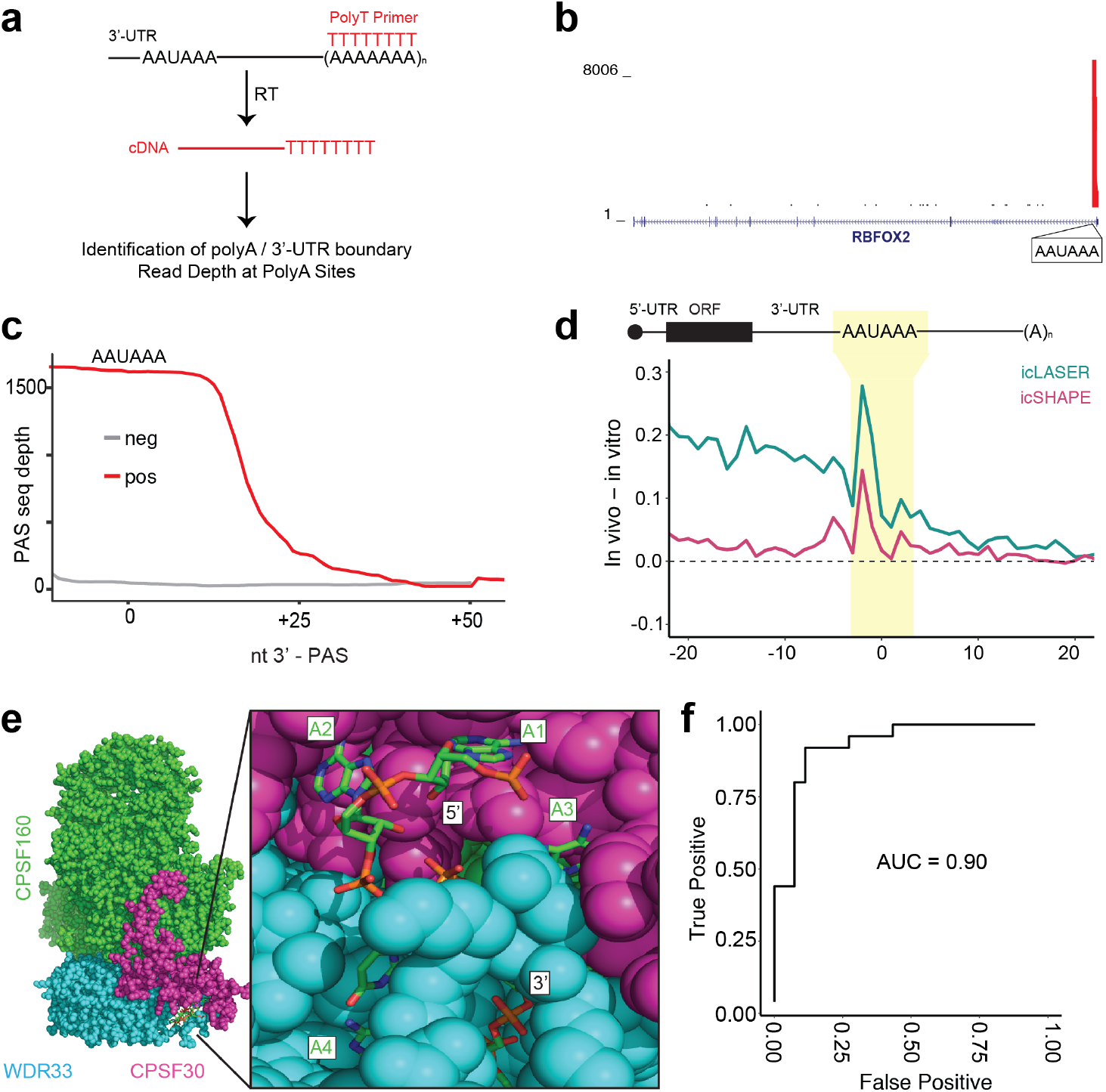
Utilizing icSHAPE and icLASER data to predict RNA polyA sites transcriptome-wide. **a.** Schematic of polyadenylation sequencing (PASseq). **b.** Genome browser track showing read density near the 3’-end of an RNA (RBFOX2) and demonstrating PAS seq specificity. **c.** Cumulative read density at PASseq-determined *in vivo* and *in vitro* polyA sites. The zero point on the X-axis is centered on the uridine residue of the AAUAAA motif. **d.** icSHAPE and icLASER data comparing in vivo and in vitro RNA structure profiling at the PAS site. **e.** Structure of the cryo-electron microscopy structure of a quaternary complex of human CPSF-160, WDR33, CPSF-30, and an AAUAAA RNA (PDB 6BLL). **f.** ROC analysis for predicting polyA sites using structure probing. The classifier was trained using icSHAPE and icLASER profiles.

## Discussion

Here, by developing novel RNA structure bi-functional probes, we extend the utility and flexibility of our previously reported RNA solvent accessibility probe, NAz. The new reagent, NAz-N3, similarly relies on specific photoactivation of an aroyl azide by long-wavelength UV light, but can subsequently be ligated to biotin using copper-free “click” chemistry, for enrichment of modified sites of adduct formation. Using this chemistry, we measured RNA solvent accessibility in K562 cells, transcriptome-wide.

With the development of transcriptome-wide RNA structure probing techniques, an exciting but thus far poorly explored aspect of these data has been the possibility to infer interactions between RNAs and their cellular partners. By employing a computational strategy (SVM) to combine icSHAPE and icLASER (as well as other reactivity-based measurements) we take a critical step towards learning the potential of these methods in predicting RBP binding and other functional RNA processing activities like PAS selection. We demonstrate the power of this approach by predicting RNA-protein binding sites for a large number of RNA binding proteins. We demonstrate that such an approach could be very powerful for measuring RNA-protein interfaces and is highly complementary with protein-centric methods such as eCLIP. Further, we utilize RNA structure probing to identify a structure signature associated with polyadenylation sites, which is due to the presence of a protein bound at the PAS site. This extension also enables structure probing to be utilized for other aspects of RNA biology, such as RNA processing.

The complexity of nucleic acid polymer structure is now well appreciated for DNA and the folding of the genome; importantly, measuring the structure of such nucleic acids and how proteins interact with DNA has been incredibly valuable for understanding how the genome is regulated to control biological processes within cells. Our goal herein is to make headway toward this goal for RNA, by the continued development of tools for transcriptome-wide measurement of RNA structure and RNA interactions that can contribute to its biological function and regulation. We anticipate that icLASER (a new aspect of transcriptome-wide RNA structure probing via solvent accessibility) will become an increasingly useful chemical tool to probe RNA structure in living cells transcriptome-wide. By demonstrating the predictive power of these tools, we expect RNA structure probing to expand its value into aspects of RNA biology.

## Contributions

D.C. worked on generating biochemical data for structure probing and libraries for RNA sequencing. C.F. worked on chemical synthesis and biochemical characterization of all compounds. W.E. worked on data analysis and RNA structure analysis. R.F. helped with generating RNA sequencing libraries. D.W. and A.M. worked on data analysis and discussions on RNA-protein interactions. X.W. and Y.S. worked to help with generating PAS seq data and WE analyzed the PAS seq datasets.

## Supporting information

Updated Supplementary Information

## Data availability

icLASER and icSHAPE datasets are deposited on GEO under the accession number GSE132099.

Additionally, icSHAPE datasets are deposited on the ENCODE database under experiments ENCSR976RFC, ENCSR803XFA, ENCSR286LXS, ENCSR992XHC, ENCSR052BBY, and ENCSR836VQU

PAS sequencing datasets are deposited on GEO under the accession number GSE145400.

## Acknowledgements

We thank the Spitale lab for their careful reading of the manuscript. This research was supported by the NIH (5UM1HG009443-03 to RCS). RCS is a Pew Biomedical Scholar.

## Online Methods

All methods pertaining to the synthesis of all probes is presented in the Supplementary Information.

### Primers used for Reverse Transcription

SAM-I (5’, ATTTAGGTGACACTATAGTT, 3’)

18s Primer 1 (5’, CCAATTACAGGGCCTCGAAA, 3’)

### SAM-I Construct (Control RNA for analysis of NAz-N_3_ structure probing)

A 94 nucleotide construct consisting of the sequence for the SAM-I riboswitch from the metF-metH2 operon of *T. tencongensis* was designed into a plasmid with IDT. The plasmid encoding the SAM-I sequence was transformed into a One Shot Top 10 chemically competent cells (ThermoFisher) and plated on lysogeny broth (LB, ThermoFischer) supplemented with 100 mg/mL ampicillin (VWR) agar plates. A single colony was selected in a 3 mL culture and grown overnight. The resulting plasmid was isolated using a QIAprep Miniprep kit (Qiagen). Transcription template was prepared by PCR in 50 μL volumes using primers directed against the T7 promoter (5’, TAATACGACTCACTATAGGG, 3’) and an adaptor sequence for reverse transcription (5’, ATTTAGGTGACACTATAGTT, 3’) with Phusion High-Fidelity PCR master mix (NEB). The mixture was treated with 1 μL of DPN1 (NEB) restriction enzyme and incubated at 37 °C for 30 min. The resulting PCR product was purified using a QIAquick PCR purification kit (Qiagen). *In vitro* transcription of SAM-I RNA was performed in 20 μL reactions using 800 ng of template SAMI DNA with a T7 Ribomax transcription kit (Promega). Samples were treated with 1 μL of RQ1 DNAse (Promega) at 37 °C for 30 min. The resulting solution was brought up to 100 μL and precipitated with 10 μL of 3 M NaOAc (pH 5.2), 1 μL glycoblue (ThermoFisher), and 300 μL ethanol. The resulting RNA stored at −80 °C for 30 minutes, centrifuged at 15,000 rpm for 15 minutes, and resuspended in 50 μL of RNase-free water. An aliquot of SAM-I was run on a denaturing PAGE gel (15% polyacrylamide, 0.5X TBE, 7 M urea) alongside a 100 bp RNA ladder (Invitrogen). The band of interest was visualized with 1x SYBR-gold stain (ThermoFisher) in water for 15 minutes. Resulting concentrations of RNA was quantified by integrating intensity of the ladder with the RNA band of interest.

### ^32^P End labeling for reverse transcription

Primer DNA was 5’ end labeled in 10 μL reactions in a T4 PNK mix (1 μL 10x T4PNK reaction buffer, 2 μL 100 mM DNA primer, 5 μL nuclease free water, 1 μL γ-^32^P-ATP, 1 μL T4 PNK, NEB). The reaction was allowed to proceed for two hours at 37 °C. Reactions were stopped by the addition of 5 μL of Gel Loading Buffer II. The reaction was loaded onto a 15% denaturing PAGE gel. The band of interest was visualized by a phosphorimager (Typhoon, GE healthcare). The resulting band was excised and eluted overnight in 400 μL of 300 mM KCl. Resulting solution was EtOH precipitated and dissolved to 8,000 counts per minute (cpm)/μL for further use in reverse transcription.

### Modification of RNA *in vitro*

5 μg total RNA (isolated from HeLa cells) or 10 pmol of *in vitro* transcribed RNA in 6 μL metal-free water was heated for 2 minutes at 95 °C. The RNA was then flash cooled on ice for 1 minute, and brought to room temperature. 3 μL of 3x RNA folding buffer (333 mM HEPES, pH 8.0, 20 mM MgCl2 and 333 mM NaCl) was added, and the RNA was allowed to equilibrate at 37 °C for 5 min. To this mixture 1 μL of 3 M NAz, 1 M NAz-N_3_ in DMSO (+) or DMSO (-) was added. Reactions were then exposed to 20 Watt lamp (Zilla Desert UVB 50) UV light for 3 minutes for NAz, and 10 minutes for NAz-N_3_. Reactions were brought up to 200 μL water, and extracted once with 200 μL acid phenol/chloroform/isoamyl alcohol (pH 4.5, Ambion), and washed twice with 200 μL chloroform (Sigma). Samples were precipitated by adding 20 μL 3 M NaOAc (pH 5.2), 1 μL glycoblue (20 μg/μL) and 600 μL EtOH. Pellets were washed twice with 70% cold ethanol and resuspended in 5 μL nuclease free water.

### Copper free click Chemistry of Modified RNA

In a reaction volume of 50 μL, modified RNA (10 pmol) was incubated with 5 μL Dibenzocyclooctyne-PEG4-biotin (DBCO-Biotin, 10 mM, Sigma) for 1 h at 37 °C in 1x PBS. Reactions were brought up to 200 μL and extracted once with 200 μL acid phenol/chloroform/isoamyl alcohol (pH 4.5), and washed twice with 200 μL chloroform. Samples were precipitated by adding 20 μL of 3 M NaOAc (pH 5.2), 1 μL glycoblue (20 μg/μL) and 600 μL EtOH. Pellets were washed twice with 70% cold ethanol and resuspended in 2 μL nuclease free water for dot blot analysis, or 10 μL nuclease free water for streptavidin enrichment assays.

### Dot blot Analysis of Enriched modified RNA

Hybond N+ membranes (GE) were pre-incubated in 10X SSC. Precipitated biotinylated total RNA was dissolved in 2 μL of RNase free water. RNA was loaded onto the Hybond membrane and crosslinked using 254 nm ultraviolet light. The membrane was incubated with blocking solution (120 mM NaCl, 16 mM Na2HPO4, 8 mM NaH2PO4, 170 mM SDS) for 30 min. To the membrane was added 1 μL Pierce high sensitivity streptavidin-HRP(ThermoFisher) in blocking solution. The membrane was washed twice with wash buffer A (1:10 blocking solution) for 30 min, and twice with wash buffer B (100 mM Tris pH 9.5, 100 mM NaCl, 20 mM MgCl2) for 5 min. Membrane was incubated with Pierce Western blotting substrate and visualized on the ChemiDoc (Biorad) under chemiluminesence hi sensitivity. To visualize RNA, the membrane was stained with a methylene blue solution (0.2% w/v methylene blue, 0.4 M sodium acetate).

### Modification of RNA in cells

HeLa cells were grown in DMEM (4.5 g/L glucose & L-glutamine [-] sodium pyruvate, Thermofisher) culture medium supplemented with 10% FBS (SAFC) and 1% penicillin streptomycin (Life technologies). K562 cells were grown in RPMI 1640 supplemented with 10% FBS, and 1% penicillin streptomycin. Cells were washed three times with Dulbecco’s phosphate-buffered saline (DPBS, Genesee) and centrifuged at 1000 RPM for 5 min. Cells (~3-6 x 10^7^) were resuspended in 45 μL DPBS. 5 μL DMSO (-), 10% final concentration, 5 μL 3 M NAz or 1 M NAz-N3 in DMSO (+) was added to the desired final concentration. Cell suspensions were subjected to UV light for 5 minutes using NAz and 10 minutes with NAz-N_3_. Cells were pelleted by centrifugation at 1000 RPM for 5 min and resuspended in 1 mL Trizol Reagent (ThermoFisher). RNA was harvested using Trizol Reagent following the manufacturer’s instructions. 500 μL of aqueous phase was then precipitated with 500 μL isopropyl alcohol at room temperature for 10 minutes. Samples were centrifugated at 15,000 RPM at 4 °C and washed twice with cold 70% EtOH.

### Enrichment of Modified RNA

In 700 μL reaction volume, 50 pmol of biotinylated RNA was added with 50 μL of prewashed Dynabeads MyOne C1 beads (ThermoFisher). The solution was then mixed at room temperature for 1 h. The beads were collected on a magnetic plate and flowthrough was saved. The beads were then washed three times with 700 μL of Biotin Wash buffer (10 mM Tris-HCl, pH 7.0, 1 mM EDTA, 4 M NaCl, 0.2% Tween). The first wash was saved and combined with the flowthrough for further analysis. Samples were later washed twice with RNase-free water. NAz-N_3_ adducts underwent harsher wash conditions was incubated twice with 700 μL Biotin wash buffer for 5 min along with two washes with RNase-free water at 70 °C. Samples were eluted twice with 44 μL formamide, 1 μL of 0.5 M EDTA, and 5 μL of 50 mM of free D-Biotin at 95 °C for 5 min. Eluted samples were diluted with 600 μL RNase-free water. All samples were purified using RNA Clean and Concentrator Kit (Zymo). Samples were eluted in 6 μL of RNase-free water and used for subsequent reverse transcription.

### Reverse transcription of modified RNA (in vitro and in vivo)

^32^P-end-labeled DNA primers were annealed to modified RNA by incubating 95 °C for two minutes, then 25 °C for two minutes, and 4 °C for 2 minutes. To the reaction, 1 μL of 5x First strand buffer, 0.5 μL nuclease free water, 0.5 μL 100 mM DTT, and 0.5 μL 10 mM dNTP’s were added. The reaction was preincubated at 52 °C for 1 min, then 1 μL superscript III (ThermoFisher) was added. Extensions were performed for 15 min. To the reaction, 1 μL 4 M sodium hydroxide was added and allowed to react for 5 min at 95 °C. The resulting complementary DNA (cDNA) was snapped cooled on ice, and ethanol precipitated according to above procedures. Purified cDNA was resuspended in 2 μL of nuclease-free water and 2 μL of Gel Loading Buffer II. cDNA products were resolved on 10% denaturing polyacrylamide gel, and visualized by a gel imager (Typhoon, GE healthcare).

### Generation of PolyA sequencing libraries

10 μg total RNA extracted with Trizol (Ambion) was fragmented with fragmentation reagent (Ambion) at 70°C for 10 minutes followed by precipitation with ethanol. Reverse transcription was performed with PASSEQ7-2 RT oligo:

[phos]NNNNAGATCGGAAGAGCGTCGTGTTCGGATCCATTAGGATCCGAGACGTGTGCTCTTCCGATCTTTTTTTTTTTTTTTTTTTT[V-Q]

and Superscript III (Invitrogen). cDNA was recovered by ethanol precipitation and 12O-2OO nucleotides of cDNA was gel-purified from 8% Urea-PAGE. Recovered cDNA was circularized with Circligase™ II (Epicentre) at 60°C overnight. Buffer E (Promega) was added in cDNA and heated at 95 °C for 2 minutes, and then cool to 37 °C slowly. Circularized cDNA was linearized by BamH I (Promega). cDNA was collected by centrifugation after ethanol precipitation. PCR was carried out with primers PE1.0 and PE2.0 containing index (Illumina). Around 200 bp of PCR products was gel-purified and submitted for sequencing (single read 100 nucleotides).

### Data analysis for icSHAPE and icLASER

Raw sequence reads were quality-checked using FastQC (29) and demultiplexed. Reads were aligned to the ENSEMBL Release 88 GRCh38 transcriptome (30)and per-position enrichment scores were calculated using the icSHAPE pipeline(21). Default parameters were used with the exception of the final filtering step, where minimum values for hit coverage and background base density were removed (-T 0 -t 0).

Publicly available eCLIP peak data for five RNA binding proteins in K562 cells was downloaded from the ENCODE project(31). Known 5bp binding motifs BiorXiv https://doi.org/10.1101/179648)) for each protein were located in each peak; peaks lacking the motif were discarded. icLASER enrichment scores were extracted for each motif position, plus five basepairs up- and downstream from the motif. Negative control sites were identified as occurrences of the same 5bp motif that fall outside of eCLIP peaks. As with binding sites, icLASER enrichment scores were extracted for a 15bp range centered on the motif. A number of negative sites equal to the number of positive sites was selected randomly from the pool of possible negatives.

icLASER analysis at poly-A signal sequences was conducted similarly; however, Ensembl-annotated polyadenylation signal sequences were used to identify positive sites, and enrichment scores were retrieved for a range of 20bp up- and downstream of the 6bp polyadenylation signal motif.

### SVM analysis of icSHAPE/icLASER and RNA-binding proteins

SVMs were implemented using libSVM (21). Both training and test data were scaled to a 0-1 scale and SVM parameters were selected using 5-fold cross-validation. For each motif, both *in vitro* and *in vivo* data sets were divided in half at random; half for the data was used to train the SVM, while the remaining half was used to test its predictions. icLASER scores at each position in the 15bp range surrounding the motif of interest were input as features. Ranges containing null values for icSHAPE or icLASER enrichment scores were discarded.

### SVM analysis of icSHAPE/icLASER and polyadenylation sites

SVM analysis was conducted as with RNA-binding proteins. icSHAPE or icLASER scores for a 46bp range centered on the polyadenylation signal sequence were used as features.

