## Supplementary material for "Transcriptome-Wide Combinatorial RNA Structure Probing in Living Cells": Updated Supplementary Information

(1) Department of Pharmaceutical Sciences, University of California, Irvine. Irvine, California. 92697, (2) Department of Developmental and Cellular Biology, University of California, Irvine. Irvine, California. 92697 (3) Department of Chemistry, Stanford University, Stanford CA 94305 (4) Department Microbiology and Molecular Genetics, University of California, Irvine. Irvine, California. 92697 (5) Department of Chemistry, University of California, Irvine. Irvine, California. 92697

($) These authors contributed equally to this manuscript.

### General

All reagents were purchased from commercial suppliers and were of analytical grade and used without further purification unless otherwise noted. 2'-azidoadenosine was purchased from TriLink BioTechnologies. Reaction progress was monitored by thin-layer chromatography on EMD 60 F254 plates, visualized with UV light, iodine, ninhydrin, $KMnO_4$, $FeCl_3$, p-anisaldehyde, 2,4-DNP, and bromocresol green stains. Compounds were purified via flash column chromatography using Sorbent Technologies 60 Å 230 x 400 mesh silica gel. Anhydrous solvents acetonitrile (MeCN), dichloromethane (DCM), methanol (MeOH), tetrahydrofuran (THF), dimethylformamide (DMF) were degassed and dried over molecular sieves. Acetone was dried over $MgSO_4$. All reaction vessels were flame dried prior to use. NMR spectra were acquired with Bruker Advanced spectrometers. All spectra were acquired at 298 K. $^1$H-NMR spectra were acquired at 400 MHz and 500 MHz. $^{13}$C-NMR spectra were acquired at 125 MHz. Chemical shifts are reported in ppm relative to residual non-deuterated NMR solvent, and coupling constants (J) are provided in Hz. All NMR spectra were analyzed using MestreNova software. Low and high-resolution electrospray ionization (ESI) mass spectra and Gas Chromatography mass spectra were collected at the University of California-Irvine Mass Spectrometry Facility. IR spectra were acquired from neat samples, unless otherwise noted, with a PerkinElmer Spectrum Two IR Spectrometer.

### Chemical Synthesis (Synthesis of NAz-$N_3$).

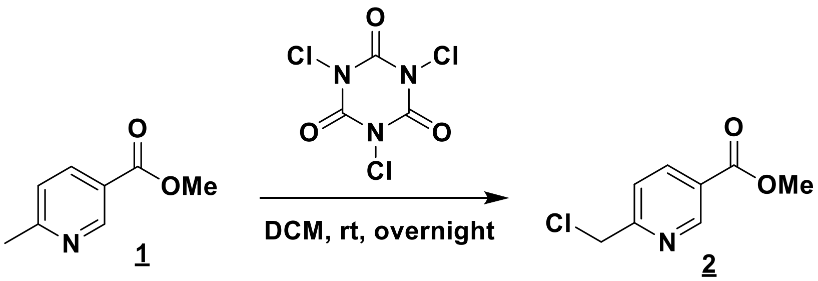


*Methyl 6-(chloromethyl)nicotinate* (**2**). To an ice-cooled solution of methyl 6-methylnicotinate (**1**) (2.00 g, 13.2 mmol) in dry DCM (10 mL) wad added 1,3,5-trichloro-1,3,5-triazinane-2,4,6-trione (3.38 g, 14.6 mmol). The resulting reaction mixture was stirred at room temperature overnight and then quenched with saturated $NaHCO_3$ (aq), extracted with DCM (20 mLx2), dried over $MgSO_4$ and concentrated under reduced pressure to afford a residue, which was purified by flash chromatography on silica gel (Hex/EtOAc) to afford **2** as a white solid (2.08 g, 85% yield). $^1$H NMR (500 MHz, $CDCl_3$) $\delta$ 3.97 (s, 3 H), 4.72 (s, 2 H), 7.60 (d, $J$ = 8.5 Hz, 1 H), 8.34 (dd, $J$ = 8, 2 Hz, 1 H), 9.17 (d, $J$ = 1.5 Hz, 1 H) ppm; $^{13}$C NMR (125 MHz, $CDCl_3$) $\delta$ 46.3, 52.7, 122.4, 125.5, 138.4, 150.7, 160.8, 165.5 ppm. HRMS (ESI) calcd for $C_8H_7ClO_2N$ [(M-H)$^-$] 184.0165, found 184.0165.

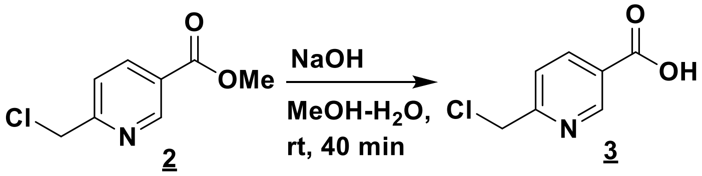


*6-(Chloromethyl)nicotinic acid (**3**)*. To an ice-cooled solution of methyl 6-(chloromethyl)nicotinate (**2**) (1.04 g, 5.40 mmol) in MeOH (5 mL) wad added a solution of NaOH (605.0 mg, 10.8 mmol) in $H_2O$ (3 mL) dropwise at room temperature. The resulting reaction solution was stirred at room temperature for 5 min and then washed with EtOAc. The resulting aqueous solution was acidified by HCl (1 N) until pH = 3, extracted with EtOAc (50 mLx3), dried over $MgSO_4$ and concentrated under reduced pressure to furnish **3** as a white solid (893.2 mg, 93% yield). $^1$H NMR (500 MHz, $d_6$-DMSO) $\delta$ 13.48 (brs, 1 H), 9.03 (s, 1 H), 8.30 (d, $J$ = 7.5 Hz, 1 H), 7.68 (d, $J$ = 8 Hz, 1 H), 4.85

(s, 2 H) ppm; $^{13}C$ NMR (125 MHz, $d_6$-DMSO) $\delta$ 46.2, 123.1, 126.0, 138.3, 150.1, 160.3, 165.9 ppm. HRMS (ESI) calcd for $C_7H_5ClNO_2$ [(M-H)$^-$] 170.0009, found 170.0006.

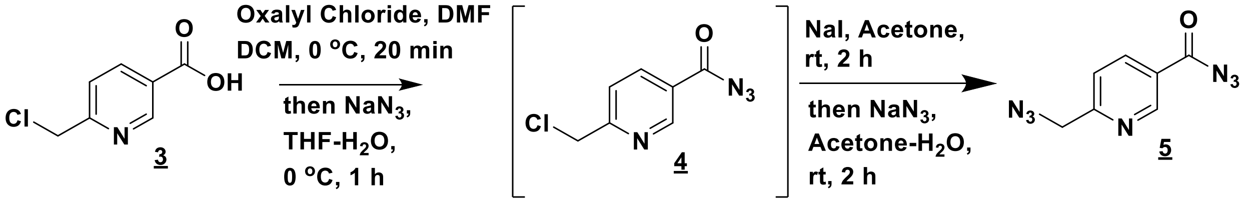


*6-(Azidomethyl)nicotinoyl azide* (**5**). To an ice-cooled suspension of 6-(chloromethyl)nicotinic acid (**3**) (506.5 mg, 2.95 mmol) and DMF (2 drops) in dry DCM (5 mL) wad added oxalyl chloride (0.38 mL, 4.43 mmol) dropwise. The resulting reaction mixture was stirred at 0 °C for 20 min and then concentrated under reduced pressure to afford a dark brown solid, which was then used for next step without further purification.

To an ice-cooled suspension of above dark brown solid in dry THF (5 mL) was added a solution of sodium azide (575.3 mg, 8.85 mmol) in water (2 mL) and the reaction mixture was then stirred at the same temperature for 2 h. The resulting reaction mixture was then quenched by concentrated HCl (0.5 mL) slowly. Saturated $NaHCO_3$ (aq.) was added to the resulting solution until pH = 8, which was then extracted with EtOAc (50 mLx3), dried over $MgSO_4$, concentrated, and purified by flash chromatography on silica gel (Hexane/EtOAc) to afford 6-(chloromethyl)nicotinoyl azide (**4**) as a pale yellow oil (352.9 mg, 61% yield from **3**). $^1H$ NMR (400 MHz, $CDCl_3$) $\delta$ 4.73 (s, 2 H), 7.62 (d, $J$ = 8.4 Hz, 1 H), 8.32 (dd, $J$ = 8.4, 2 Hz, 1 H), 9.15 (d, $J$ = 2 Hz, 1 H) ppm; $^{13}C$ NMR (125 MHz, $CDCl_3$) $\delta$ 46.1, 122.7, 126.0, 138.2, 150.4, 162.1, 171.0 ppm. HRMS (ESI): Decomposition occurred during analysis due to the nature of acyl azide.[1]

To a solution of **4** (352.9 mg, 1.80 mmol) in acetone (5 mL) was added sodium iodide (807.2 mg, 5.39 mmol) at room temperature. The resulting reaction mixture were stirred at room temperature for 3 h and then concentrated, diluted in $H_2O$, extracted with EtOAc twice, dried over $MgSO_4$, and concentrated again to afford a pale yellow solid, which was used for next step without further purification.

To a solution of above pale yellow solid in acetone (4 mL) was added a solution of sodium azide (350.4 mg, 5.39 mmol) in water (1 mL) at room temperature. The resulting reaction solution was stirred at room temperature for 1 h, concentrated under reduced pressure and diluted again in EtOAc and water. After extracted with EtOAc twice, the combined organic phase was dried over $MgSO_4$, concentrated under reduced pressure to afford a residue, which was purified by chromatography on silica gel (Hexane/EtOAc) to afford **5 (NAz-N$_3$)** as a pale yellow solid (333.0 mg, 90% yield from **4**, 56% yield from **3**). $^1H$ NMR (500 MHz, $CDCl_3$) $\delta$ 4.59 (s, 2 H), 7.48 (d, $J$ = 8 Hz, 1 H), 8.32 (dd, $J$ = 8, 2 Hz, 1 H), 9.18 (d, $J$ = 2 Hz, 1 H) ppm; $^{13}C$ NMR (125 MHz, $CDCl_3$) $\delta$ 55.5, 121.6, 125.9, 138.1, 150.7, 161.6, 171.1 ppm. HRMS (ESI): Decomposition occurred during analysis due to the nature of acyl azide.

<u>**Chemical Synthesis Spectra for Naz-$N_3$**</u>

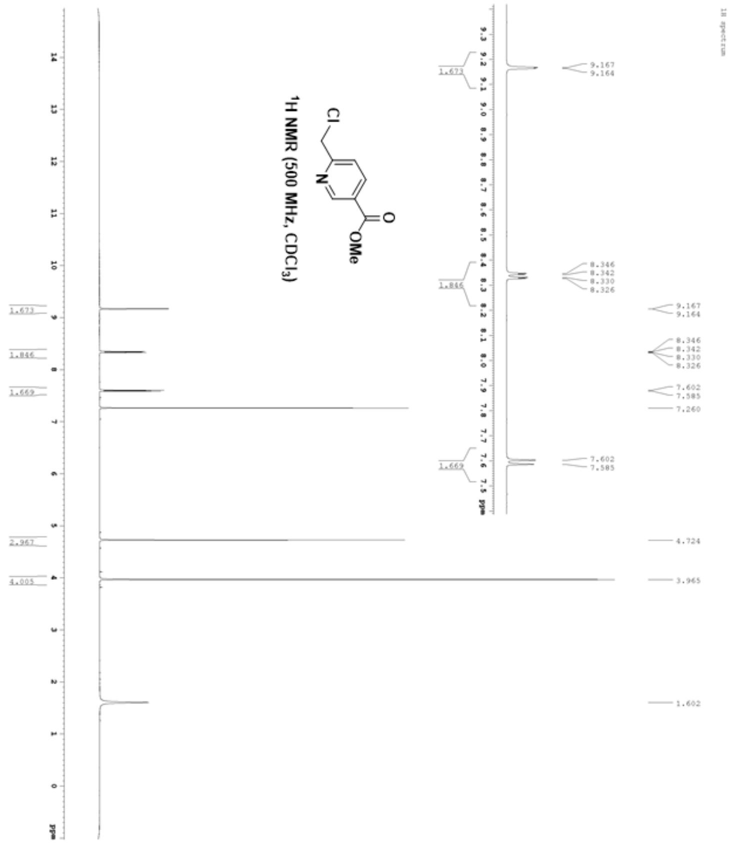

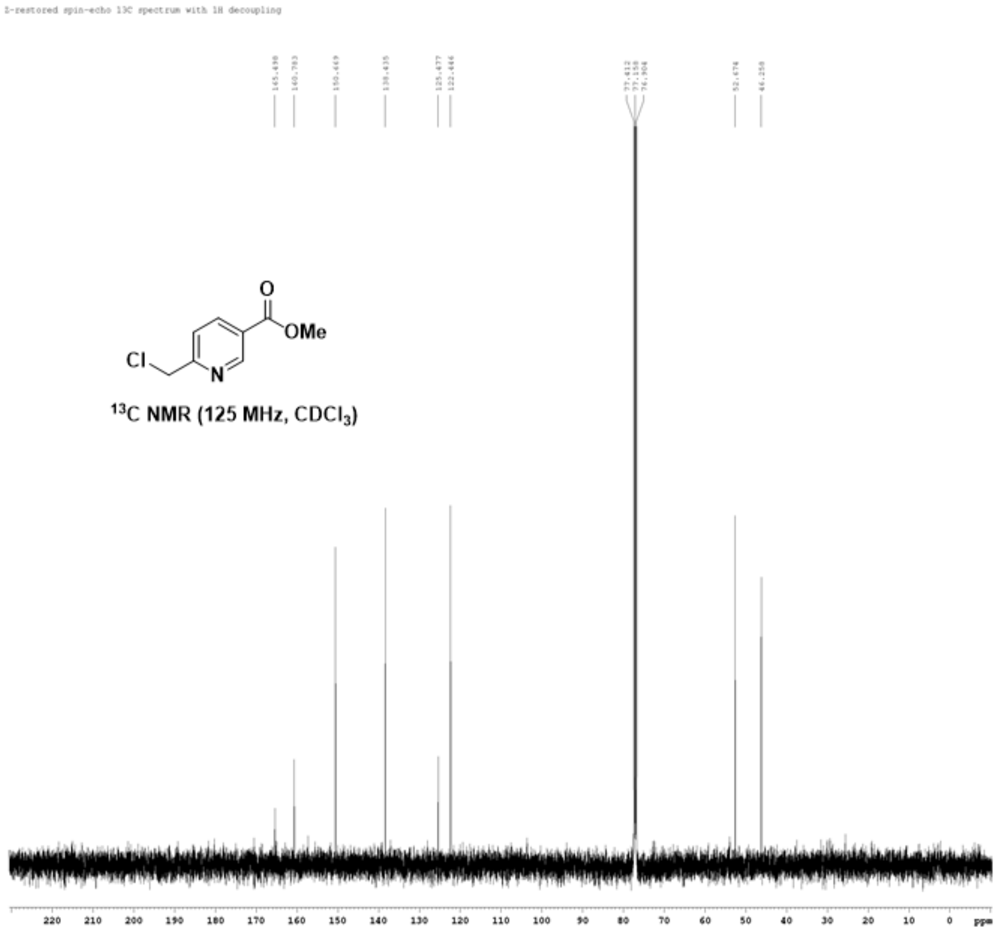


$^{13}C$ NMR (125 MHz, $CDCl_3$)

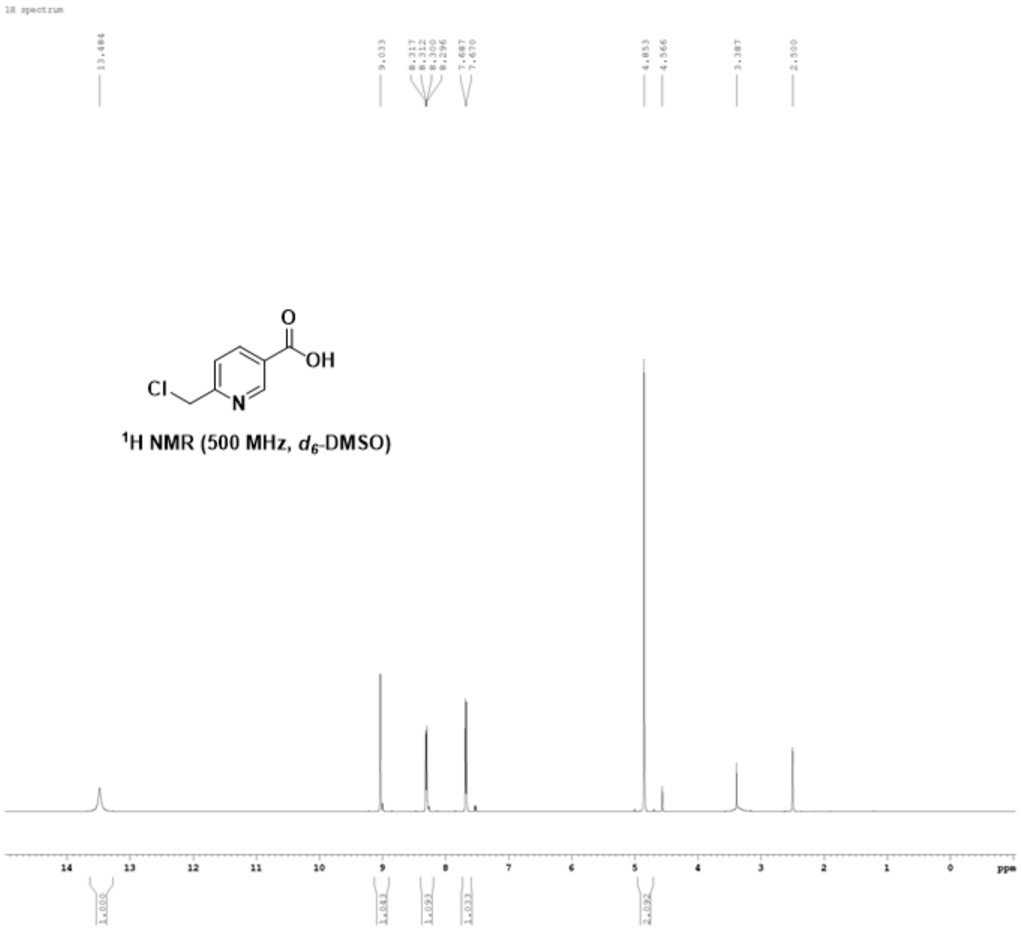
1H spectrum
13.484
9.033
8.317
8.312
8.300
8.296
7.687
7.670
4.853
4.566
3.387
2.500
Cl
N
O
OH
¹H NMR (500 MHz, d6-DMSO)
14
13
12
11
10
9
8
7
6
5
4
3
2
1
0
ppm
1.000
1.043
1.093
1.033
2.092

I-restored spin-echo 13C spectrum with 1H decoupling

165.935
160.292
150.095
138.263
126.010
123.099

46.240
40.021
39.854
39.687
39.520
39.353
39.186
39.019

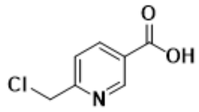

$^{13}C$ NMR (125 MHz, $d_6$-DMSO)

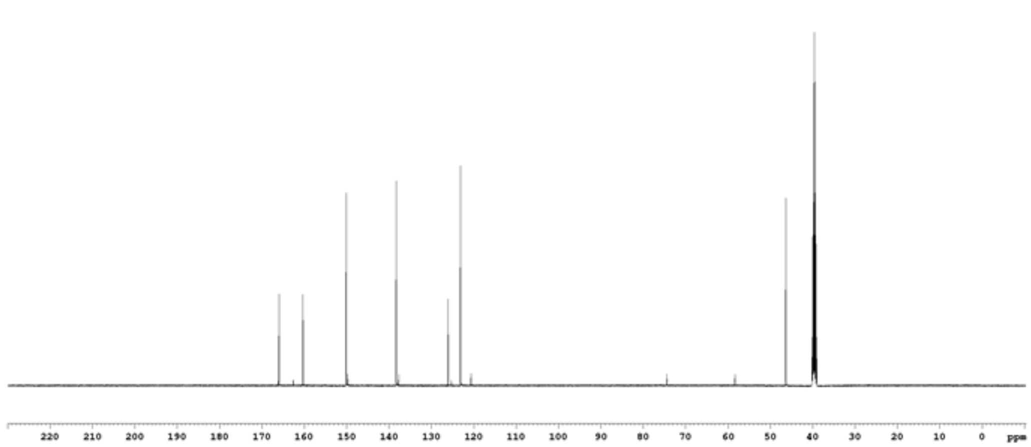

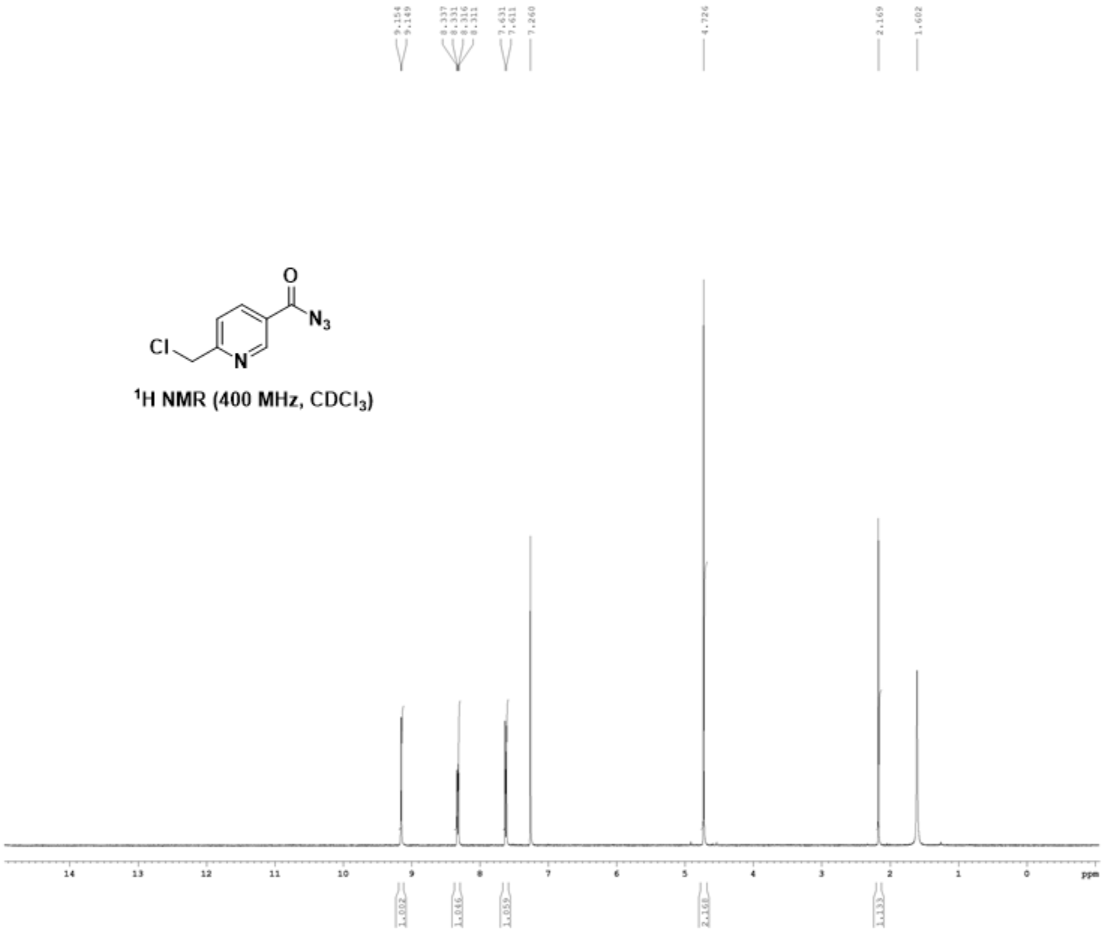

Cl
N
O
N3
1H NMR (400 MHz, CDCl3)
9.154
9.149
8.337
8.331
8.316
8.311
7.631
7.611
7.260
4.726
2.169
1.602
14
13
12
11
10
9
8
7
6
5
4
3
2
1
0
ppm
1.002
1.046
1.059
2.168
1.133

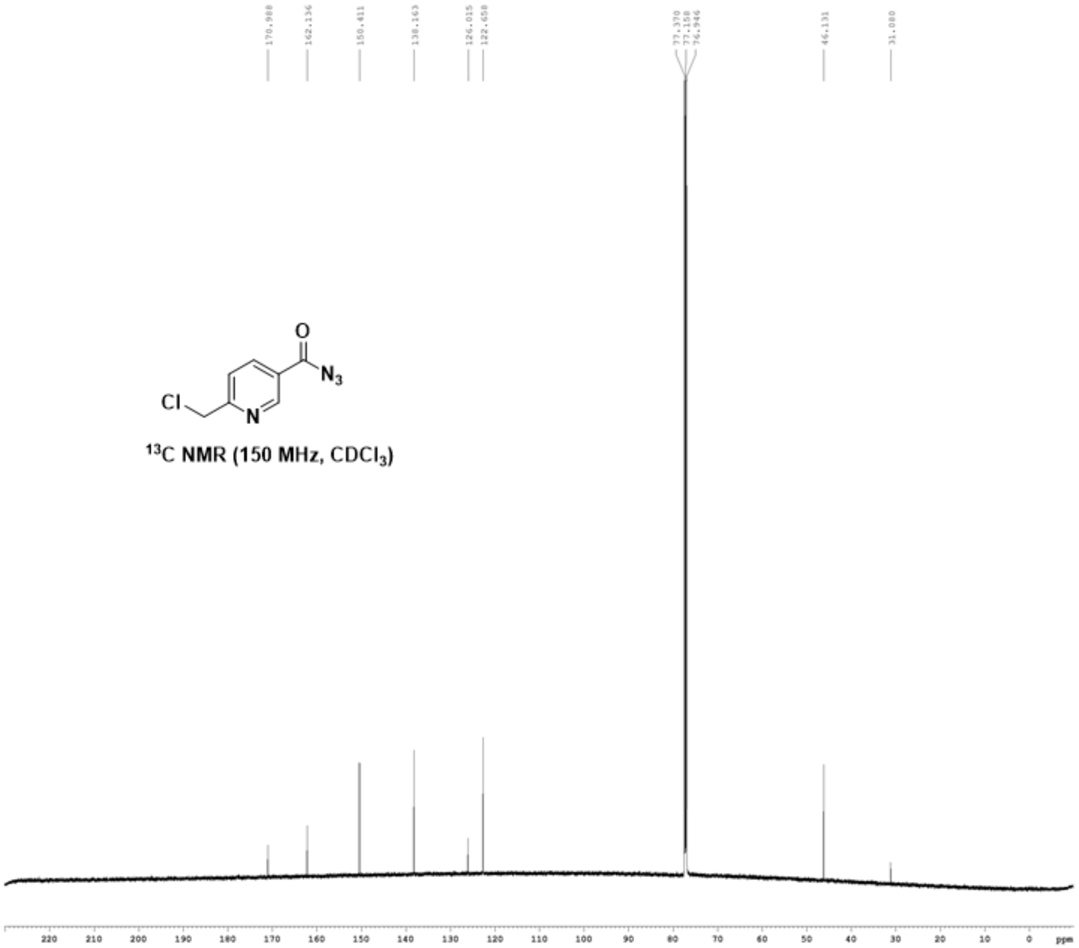

170.988
162.136
150.411
138.163
126.015
122.658
77.370
77.158
76.946
46.131
31.080
O
$N_3$
Cl
N
$^{13}C$ NMR (150 MHz, $CDCl_3$)
220 210 200 190 180 170 160 150 140 130 120 110 100 90 80 70 60 50 40 30 20 10 0 ppm

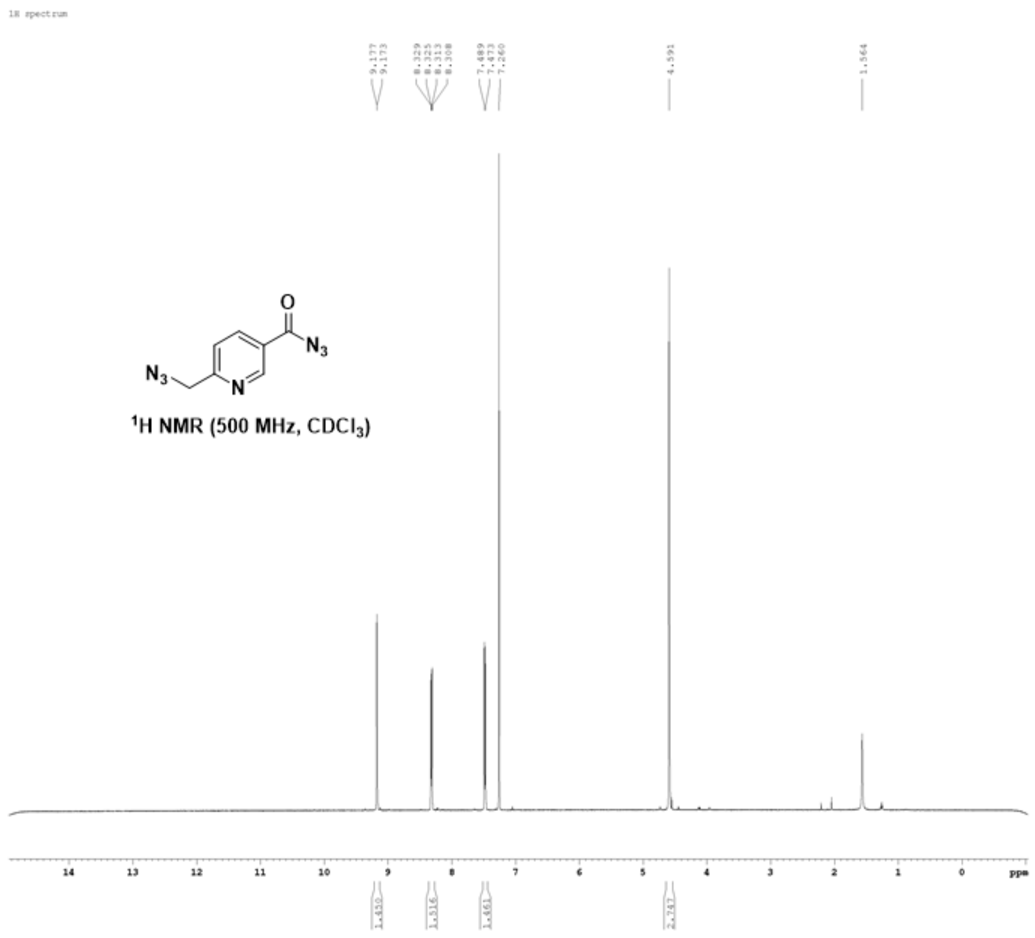

1H spectrum
9.177
9.173
8.329
8.325
8.313
8.308
7.489
7.473
7.260
4.591
1.564
N3
O
N3
N
1H NMR (500 MHz, CDCl3)
14
13
12
11
10
9
8
7
6
5
4
3
2
1
0
ppm
1.450
1.516
1.461
2.747

### Chemical Synthesis (Synthesis of Control Compounds 1 and 2 for UV-VIS studies.

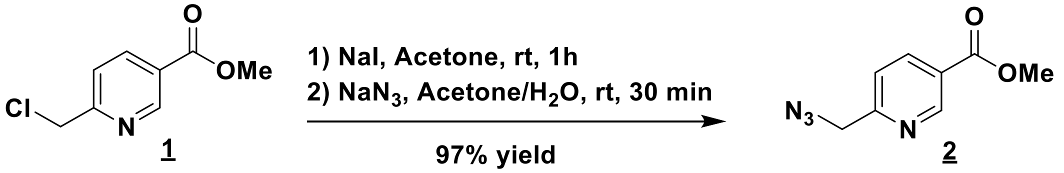


*Methyl 6-(azidomethyl)nicotinate* (**2**). To a solution of methyl 6-(chloromethyl)nicotinate (**1**) (722 mg, 3.89 mmol) in acetone (10 mL) was added sodium iodide (1.17 g, 7.78 mmol), and the resulting reaction mixture was stirred at room temperature for 1 h, followed by filtration which afforded a brown solution. A solution of $NaN_3$ (379 mg, 5.84 mmol) in water (5 mL) was added to above brown solution. After stirred at room temperature for 30 min, the reaction mixture was extracted with EtOAc twice, dried over $MgSO_4$ and concentrated under reduced pressure to afford a residue, which was purified by flash chromatography on silica gel (Hex/EtOAc) to afford **2** as a yellow solid (727 mg, 97% yield from **1**). $^{1}$H NMR (400 MHz, $CDCl_3$) $\delta$ 3.96 (s, 3 H), 4.57 (s, 2 H), 7.45 (d, $J$ = 8 Hz, 1 H), 8.33 (dd, $J$ = 8, 2 Hz, 1 H), 9.19 (d, $J$ = 2 Hz, 1 H) ppm; $^{13}$C NMR (125 MHz, $CDCl_3$) $\delta$ 52.6, 55.6, 121.5, 125.4, 138.3, 151.0, 160.2, 165.6 ppm.

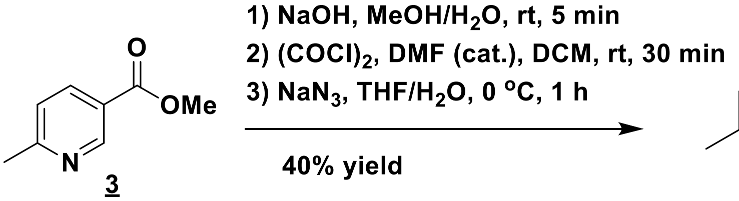


*6-Methylnicotinoyl azide (**4**)*. To a solution of methyl 6-methylnicotinate (**3**) (1.05 g, 6.95 mmol) in MeOH (10 mL) wad added a solution of NaOH (556 mg, 13.9 mmol) in $H_2O$ (1 mL) dropwise at room temperature. The resulting reaction solution was stirred at room temperature for 5 min and then neutralized with HCl (aq.) until pH = 5. The reaction mixture was concentrated directly under reduced pressure to afford a residue, which was dried under high vacuum for 2 h to afford a brown solid and used for next step without further purification. To above brown solid was added a solution of oxalyl chloride in DCM (5.20 mL, 2 M, 10.4 mmol) at room temperature, followed by a drop of DMF, and the resulting suspension was stirred at room temperature for 30 min, then concentrated under reduced pressure to furnish a residue, which was resuspended in anhydrous THF (10 mL) under argon and cooled down to 0 °C. A solution of sodium azide (1.36 g, 20.9 mmol) in water (5 mL) was added to above suspension and the reaction mixture was stirred at 0 °C over 1 h. The reaction mixture was diluted with $NaHCO_3$ (aq.), extracted with EtOAc twice, dried over $Na_2SO_4$, filtrated, concentrated under reduced pressure to afford a residue, which was purified by chromatography on silica gel to produce **4** as a pale yellow solid (446 mg, 40% yield from **3**). $^{1}$H NMR (400 MHz, $CDCl_3$) $\delta$ 2.64 (s, 3 H), 7.26 (d, $J$ = 8.4 Hz, 1 H), 8.16 (dd, $J$ = 8, 2.4 Hz, 1 H), 9.09 (d, $J$ = 2 Hz, 1 H) ppm; $^{13}$C NMR (100 MHz, $CDCl_3$) $\delta$ 25.1, 123.3, 124.1, 137.2, 150.5, 164.9, 171.5 ppm. HRMS (ESI) not obtained because of decomposition due to the nature of acyl azide.

**Chemical Synthesis (Synthesis Spectra  of Control Compounds 1 and 2 for UV-VIS studies.**

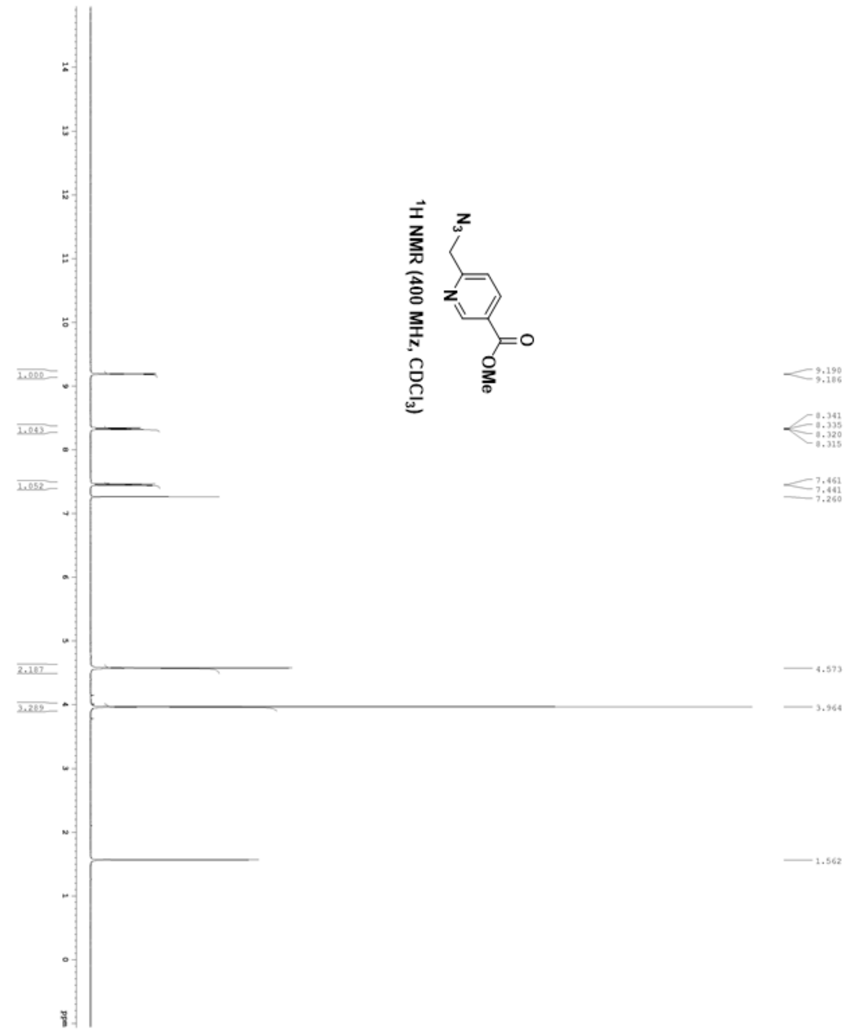

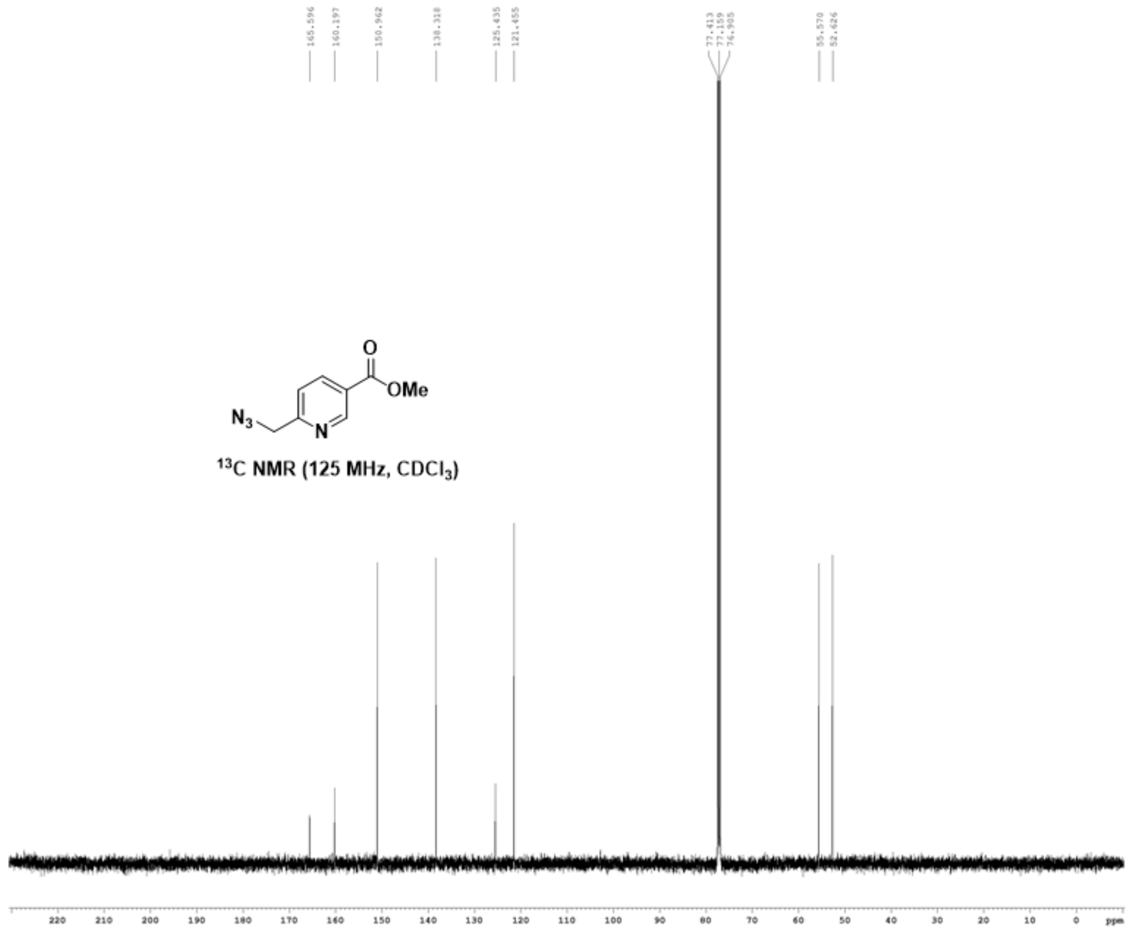

165.596
160.197
150.962
138.318
125.435
121.455
77.413
77.159
76.905
55.570
52.626
N3
N
O
OMe
13C NMR (125 MHz, CDCl3)
220
210
200
190
180
170
160
150
140
130
120
110
100
90
80
70
60
50
40
30
20
10
0
ppm

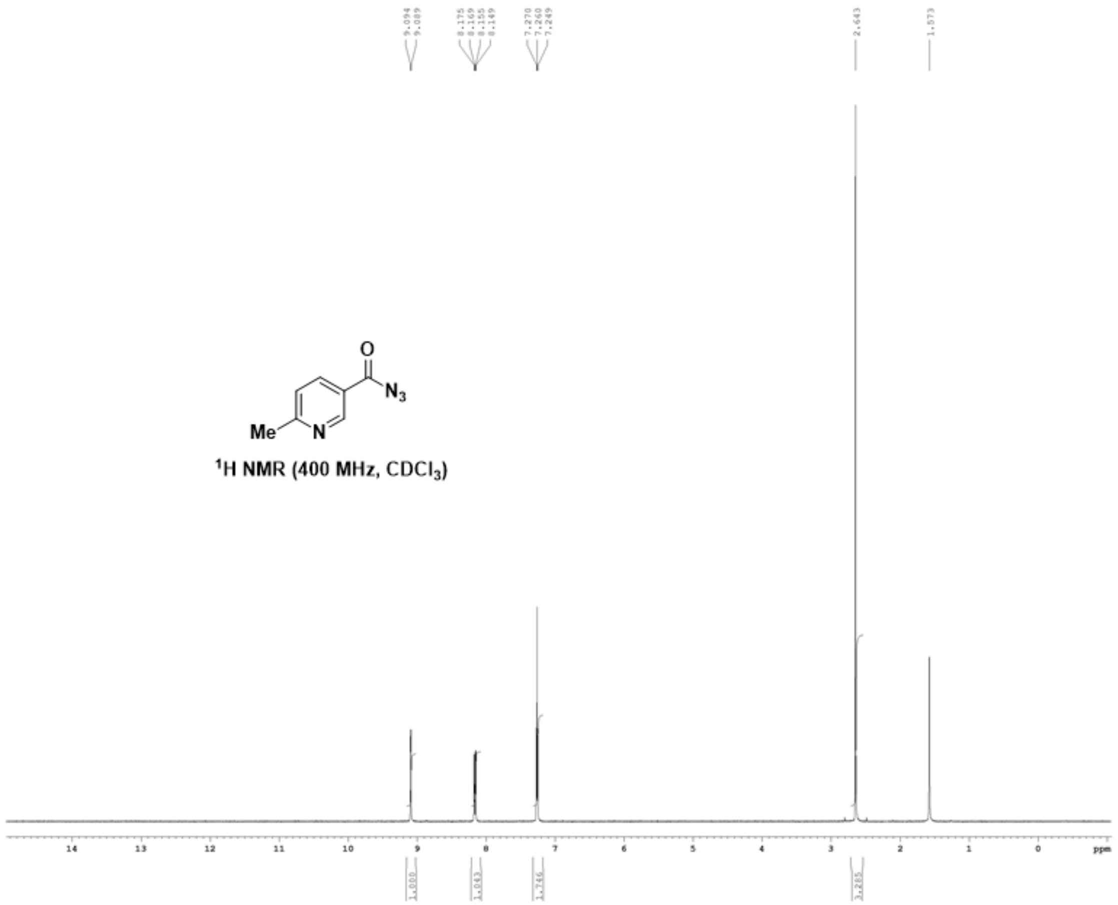

$^1$H NMR (400 MHz, $CDCl_3$)

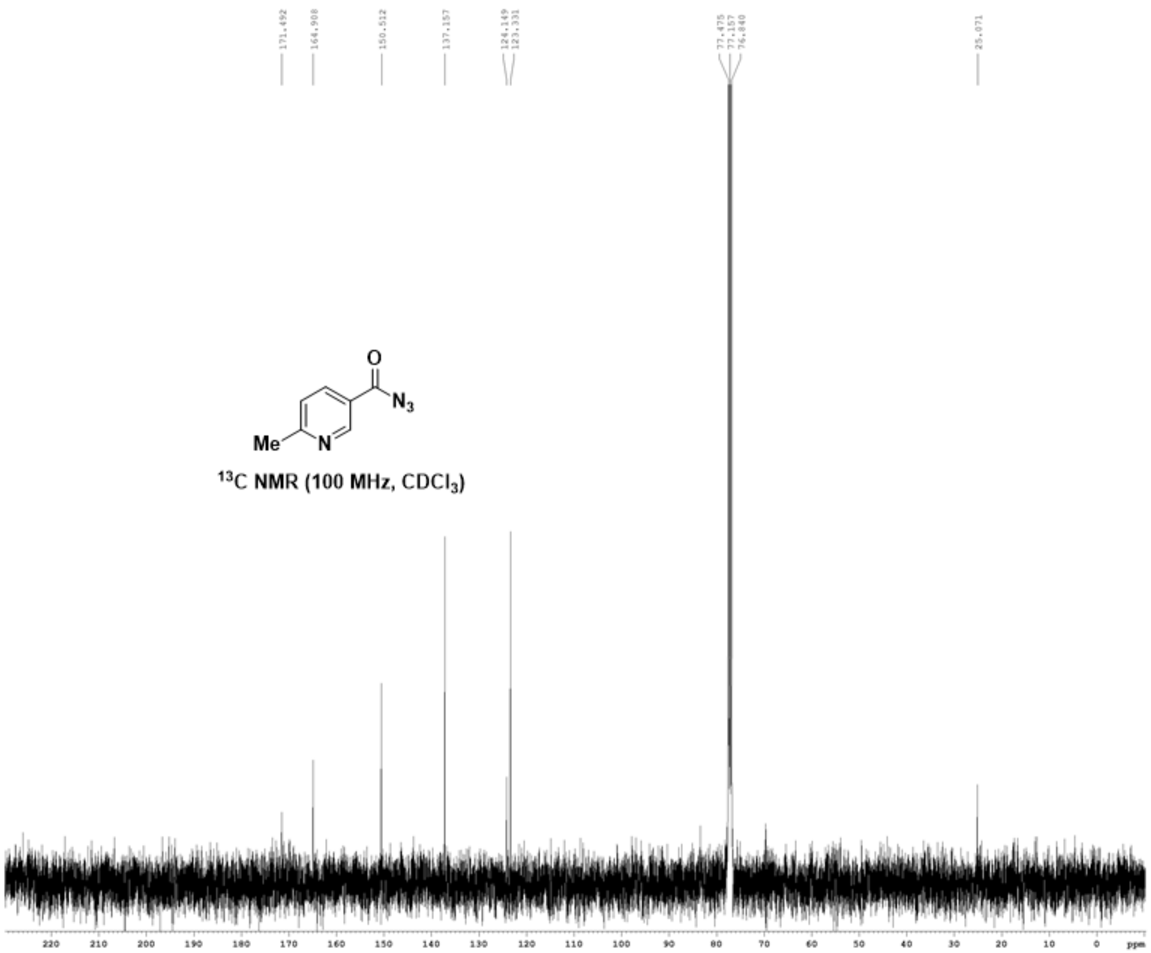

171.492
164.908
150.512
137.157
124.149
123.331
77.475
77.157
76.840
25.071
Me
N
O
N3
13C NMR (100 MHz, CDCl3)
220
210
200
190
180
170
160
150
140
130
120
110
100
90
80
70
60
50
40
30
20
10
0
ppm

**upplementary Information Figures.**

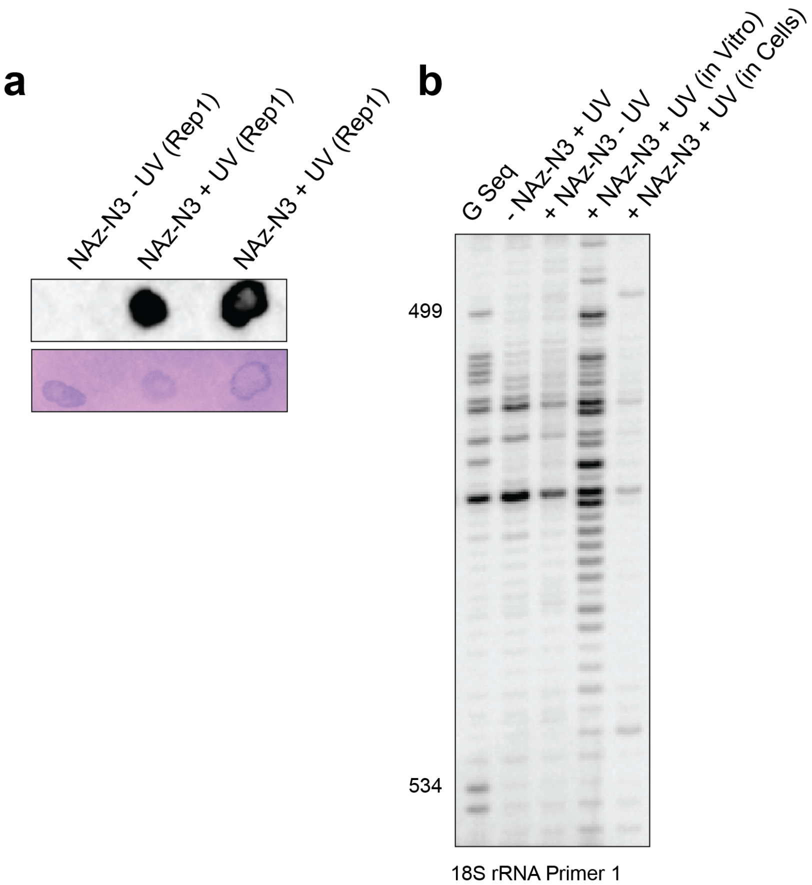


**Supplementary Information Figure 1.** NAz-$N_3$ can modify RNA in living cells. **a.** Dot blot showing biotinylation on isolated RNA after NAz-$N_3$ treatment. **b.** Reverse transcription gel demonstrating similar RNA modification pattern that has been observed with NAz.

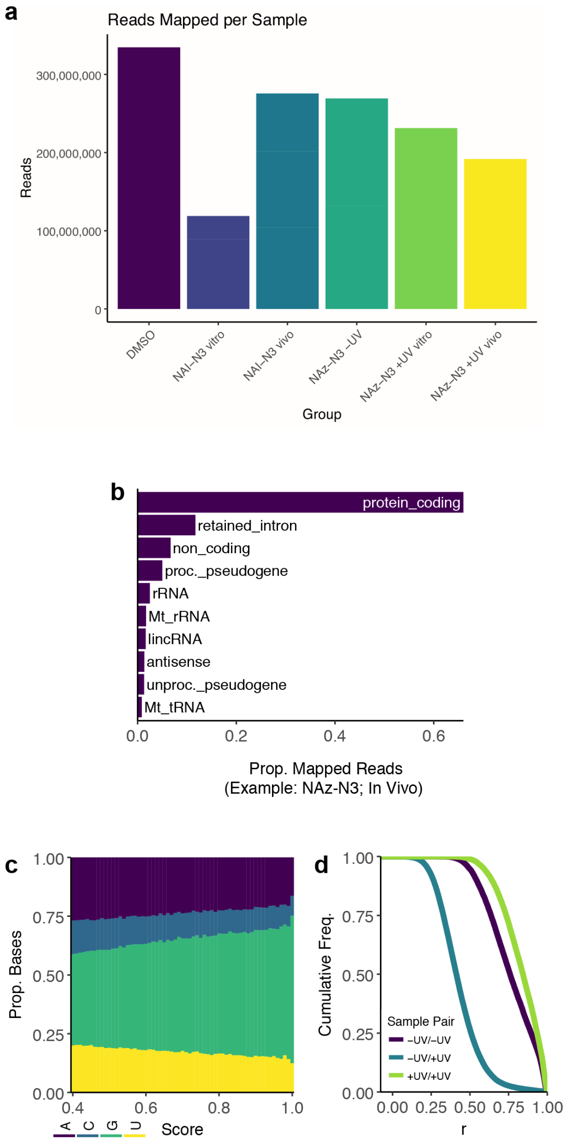


**Supplementary Information Figure 2.** Analysis of read data from each dataset. **a.** Mapped read depth of each dataset. **b.** Read distribution by RNA type. **c.** Proportion of RT-stops by nucleotide identity. **d.** Correlation between datasets.

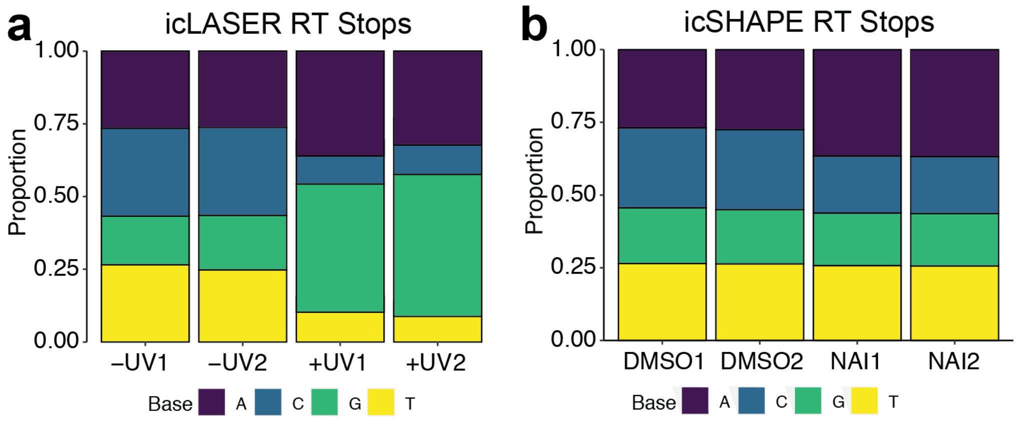


**Supplementary Information Figure 3. Plot of calculated RT-stops for icLASER and icSHAPE profiles.**

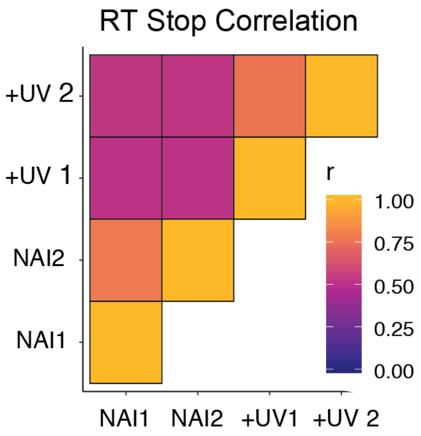


**Supplementary Figure 4. Correlation matrix of RT stops for both icSHAPE and icLASER and comparison between datasets.**

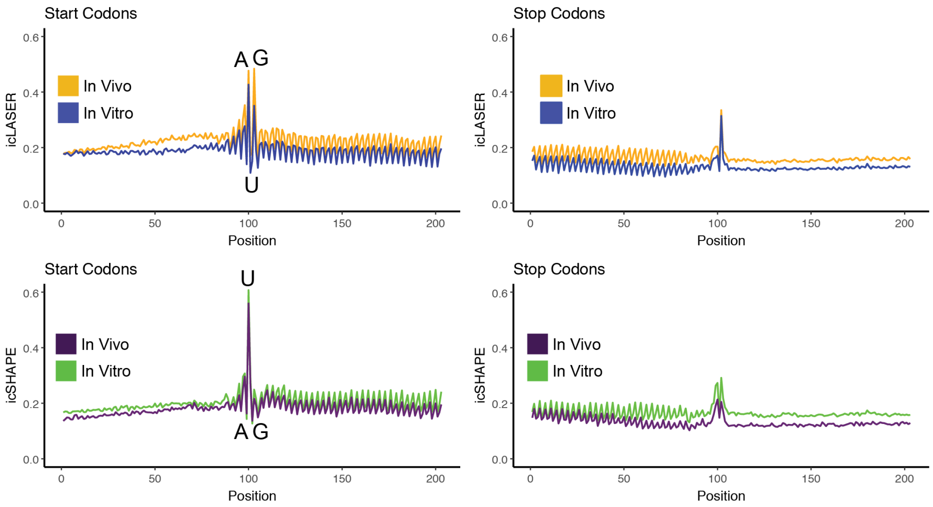


**Supplementary Information Figure 5. Comparison structure profiles for mRNAs, between icLASER and icSHAPE.**

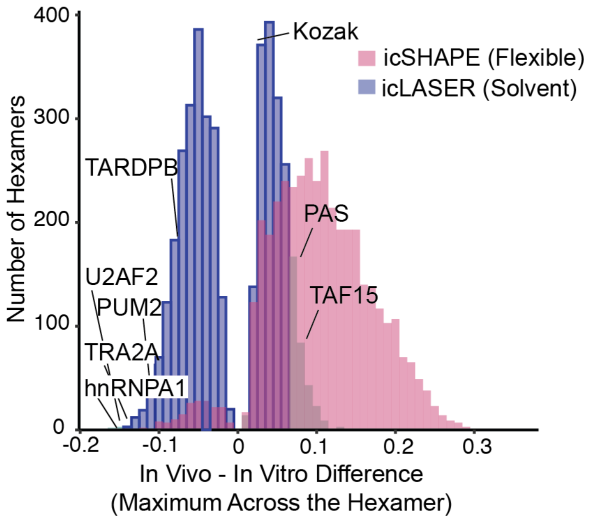


**Supplementary Figure 6. Maximum RT-stop differences comparing to structure probing in living cells and RNA refolded after RNA extraction.** Hexamer analysis between the two datasets demonstrates differences that may be due to protein binding or interactions of RNA within the unique cellular environment.
